# Implementation of a novel optogenetic tool in mammalian cells based on a split T7 RNA polymerase

**DOI:** 10.1101/2021.10.27.466068

**Authors:** Sara Dionisi, Karol Piera, Armin Baumschlager, Mustafa Khammash

## Abstract

Optogenetic tools are widely used to control gene expression dynamics both in prokaryotic and eukaryotic cells. These tools are used in a variety of biological applications from stem cell differentiation to metabolic engineering. Despite some tools already available in bacteria, no light-inducible system currently exists to orthogonally control gene expression in mammalian cells. Such a tool would be particularly important in synthetic biology, where orthogonality is advantageous to achieve robust activation of synthetic networks. Here we implement, characterize and optimize a new orthogonal optogenetic tool in mammalian cells based on a previously published system in bacteria called Opto-T7RNAPs. The tool consists of a split T7 RNA polymerase coupled with the blue light-inducible magnets system (mammalian OptoT7 – mOptoT7). In our study we exploited the T7 polymerase’s viral origins to tune our system’s expression level, reaching up to 20-fold change activation over the dark control. mOptoT7 is used here to generate mRNA for protein expression, shRNA for protein inhibition and Pepper aptamer for RNA visualization. Moreover, we show that mOptoT7 can mitigate gene expression burden when compared to other optogenetic constructs. These properties make mOptoT7 a new powerful tool to use when orthogonality and viral-like RNA species are desired in both synthetic biology and basic science applications.

## INTRODUCTION

The ability to precisely control gene expression in time and space is essential to answer many open questions in biology ranging from development to metabolic processes. Traditional studies that investigate gene expression and function mostly rely on overexpression, knockdowns or knockouts of the gene of interest ^1–3^. This, however, is done at the expense of gaining information on the expression dynamics. In recent years, the field of synthetic biology has helped to address some of these challenges with the use of small molecules regulators, offering powerful tools to control gene expression ^4^. For example, systems based on gas or food additives have been used to activate gene expression in the context of genetic circuits ^5–7^. However, these approaches are limited by slow dynamics, a lack of spatial control and burden on the cellular resources. In synthetic biology, resource allocation is one of the main problems associated with engineered genetic circuits. These gene-based networks can create substantial burden on the cellular machineries challenging their use for therapy or downstream applications ^8–10^.

Light-inducible (“optogenetic”) systems offer major advantages compared to chemical-based approaches ^11–13^. These tools allow for tight dynamics, spatial and temporal control, and can either regulate gene expression by activating/repressing genes, or by restoring protein functions. To date, many optogenetic tools are available for both bacterial and mammalian cells ^14,15^. However, in mammalian cells, the complexity of gene regulatory mechanisms can make it difficult to use these tools robustly and consistently in different cell lines or between different cell states. In particular, little is known about the way these systems induce cellular stress and how this in turn affects gene regulation. No optogenetic system is currently available to decouple synthetic networks from the cellular transcriptional and translational machineries, which would provide a potential solution to some of these problems.

An optogenetic system that can function orthogonally to the cellular machinery, should ideally be independent from the cellular resources. In bacteria, synthetic systems can be decoupled from cellular transcription regulation mechanisms by using the T7 RNA polymerase ^16–18^. This polymerase, which originates from the T7 phage, can transcribe RNA at a very high level and works completely orthogonally to the cellular machinery, making synthetic circuits that use it more robust and predictable, as suggested by Segall-Shapiro et al. and Shis and Bennett ^16,17^. Despite few attempts to use T7 RNA polymerase in mammalian cells both constitutively or induced by chemicals^19–25^, no optogenetic systems based on this polymerase are currently available.

Here we implement, characterize and optimize a new optogenetic tool in mammalian cells (mOptoT7) that is based on a previously published optogenetic system in bacteria^26^. This tool consists of a split T7 polymerase coupled to photoregulators called Magnets^27,28^, which heterodimerize upon blue light exposure and return to monomers in the dark. mOptoT7 can carry out its function both in the cytoplasm and in the nucleus (Fig. 1a) ^22^, and has the unique characteristic of being orthogonal to the cellular machinery for transcription. Furthermore, due to its viral origin, mOptoT7 generates RNA species that are not normally present in mammalian cells. By exploiting this feature, we optimize mOptoT7 expression level to reach a maximum of almost 20-fold change induction over the dark control. We demonstrate that mOptoT7 can be used to generate different responses in HEK293T cells, making it an ideal tool for applications where light induction and orthogonality are desired. In particular, we induced mRNA and shRNA production for protein expression and inhibition, respectively and Pepper RNA aptamer for RNA visualization. Finally, we showed that mOptoT7 can be used to mitigate gene expression burden compared to another optogenetic tool.

**Figure 1.**
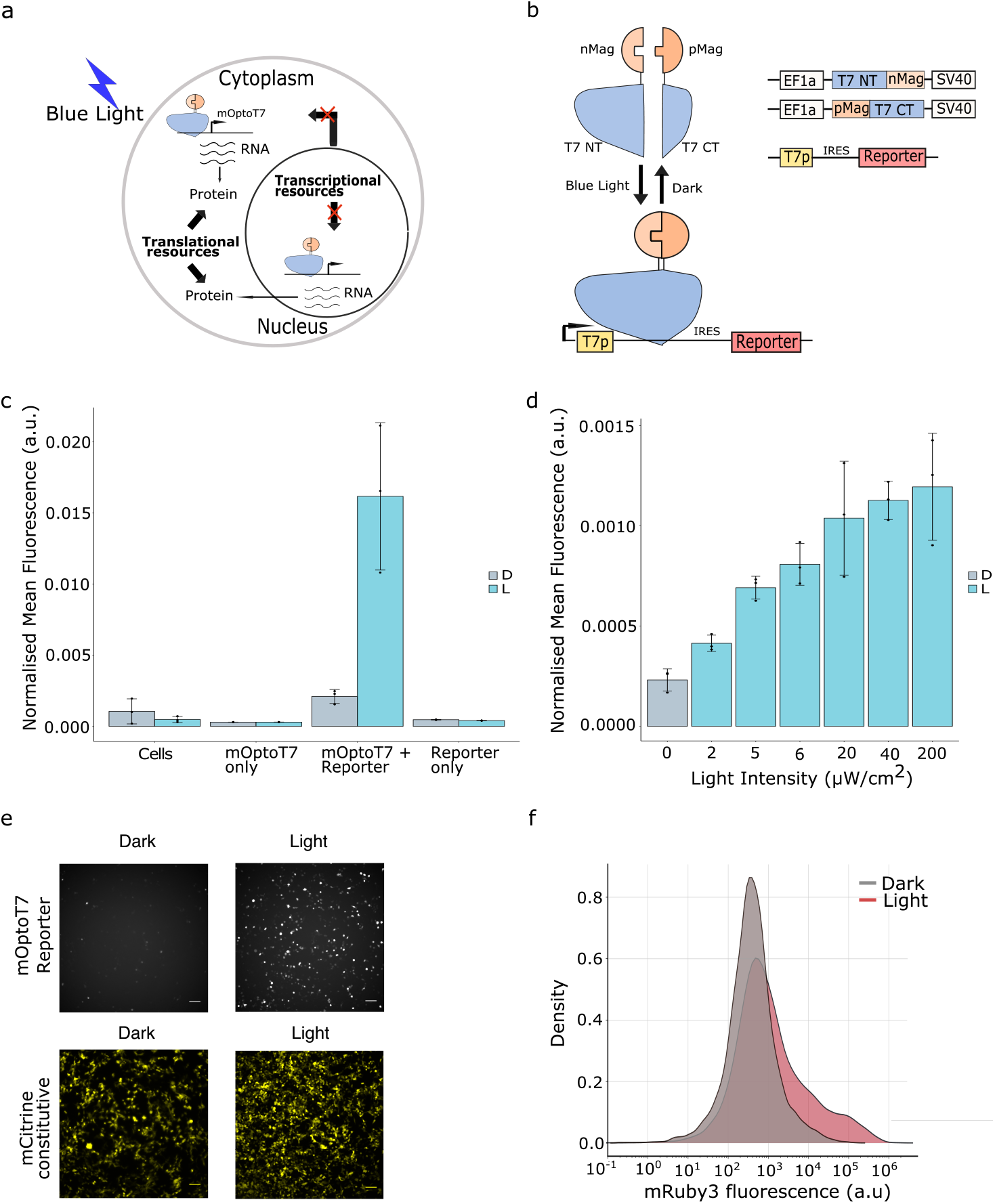
Implementation of Opto-T7RNAP in HEK293T cells. (a) Overview of mOptoT7 function in mammalin cells. (b) Experimental design. Opto-T7RNAP is transfected in HEK293T cells together with mRuby3 reporter under the control of the T7 promoter. IRES2 sequence is used to allow for translation initiation. (c) Opto-T7RNAP expression after 24h of constant blue light illumination in saturating conditions. Background fluorescence from only cells and only reporter expression is included. D=dark, L=light (d) Dose response curve of mOpto-T7RNAP with increasing light illumination. D=dark, L=light. (e) Microscopy images of mRuby3 reporter activation after 24h of constant illumination in saturating conditions compared to the dark control. mCitrine is used as constitutive color as a measure of transfection effeciency. Scale bar = 100 micron (f) Kernel density estimation plot showing mRuby3 expression in the dark vs saturating light after 24h of costant blue light illumination. Flow cytometry data are normalized to the constitutively expressed mCitrine. Saturating light: 200μW/cm2.

## MATERIAL AND METHODS

### Cell culture and Transfection

HEK293T cells (ATCC, strain number CRL-3216) were maintained in Dulbecco’s modified Eagle medium (DMEM, Gibco) supplemented with 10% FBS (Sigma-Aldrich), 1% penicillin/streptomycin, 1X GlutaMAX (Gibco) and 1mM Sodium Pyruvate (Gibco). Cells were kept at 37 °C and 5% CO_2_. Transfections were performed in a 24 well black plate (PerkinElmer) format for flow cytometry. Cells were seeded in 24 well plates at a density of 8×10^4^ cells/well one day before transfection or at 1.6×10^5^ for transfections done in suspension. HEK293T were transfected with Polyethylenimine (PEI) (Mw 40000; Polysciences, Inc.) using a ratio of 1:3 (μg DNA to μg PEI) with a total of 500 μg of DNA/well for 24 well plates. OptiMEM I reduced serum media (Gibco) was used to separately dilute both DNA and PEI. Once mixed, DNA and PEI complexes were incubated for 20 minutes at room temperature prior to addition to the cells. After transfection, cells were kept in the dark for approximately 24h before starting illumination.

### Light induction

For Flow cytometry measurements, cells were illuminated with 470nm LEDs (Super Bright LEDs Inc) using a modified version of the Light Plate Apparatus (LPA) previously described ^29^. The LPA was modified by adding a 2 cm aluminum heatsink and a ventilator both connected to the PCB in order to improve heat dissipation. In addition, a second adaptor and 2 layers of filter papers were added to allow a uniform distribution of light (Supplementary Fig.2). Cells were illuminated with constant or pulsed light (of different duration) as stated in the specific experimental conditions. The intensity of light received by the cells was measured to be 200 μW/cm2 at saturation using the S175C - Microscope Slide Thermal Power Sensor from ThorLabs. The control plate was kept in the dark during the entire experiment.

### Flow Cytometry analysis

HEK293T cells were analyzed with CytoFLEX S Flow Cytometer (Beckman Coulter) after 24h or 48h of illumination and using 488 and 561 lasers with 530/11 nm and 610/20nm OD1 bandpass filters, respectively. Prior to measurement, cells were washed once with DPBS (Thermo Fisher) and incubated with 100 μL of Accutase solution (Sigma-Aldrich) to allow detachment. For each sample, live and single cells were selected using FCS/SSC parameters and when necessary (using fluorescent proteins with overlapping emission spectra), a compensation matrix was created using single color controls and unstained cells (Supplementary Fig. 5). In every experiment, >20000 singlet events were collected for each sample and data analysis was performed using Cytoflow Software and a customized R code.

### Plasmid Construction

All plasmids were constructed using standard restriction digestion cloning or using Golden Gate assembly and a previously described ^30^ yeast toolkit (YTK) with customized parts for use in mammalian cells. In the standard cloning, PCR amplifications were performed using Phusion Flash high fidelity DNA polymerase (ThermoFisher Scientific) and ligation reactions were made using 1:3 ratios of vector plasmid:insert and incubation time of 1h at room temperature. All constructs were chemically-transformed into TOP10 *E. coli* cells and checked through sequencing (Microsynth). All relevant plasmid sequences can be found in the supplementary materials.

### Live-cell imaging

A Nikon Ti2e inverted microscope (Nikon Instruments), that is equipped with an ORCA Flash4.0 LT+ camera and a chamber for CO_2_ and temperature regulation, was used. Cells were always kept at 37 °C and with 5% CO_2_. Cells were imaged after 24h to 72h from transfection with constant illumination (performed after 24h from transfection) or after being in the dark for the same amount of time. For all experiments, mRuby3/mOrange2, mCitrine and miRFP670 fluorescence was imaged using NIS element software and with 561/4, 543/22 and 692/40 filters, respectively (BrightLine HC). CFI Plan Apochromat Lambda 10X (N.A. 0.45, W.D. 4.0mm), CFI S Plan Fluor ELWD 20XC (N.A. 0.45, W.D. 8.2-6.9mm) or CFI S Plan Fluor ELWD 40XC (N.A. 0.6, W.D. 3.6-2.8mm) were used. For brightfield imaging - LED 100 (Märzhäuser Wetzlar GmbH & Co. KG) with diffuser and green interference filter placed in the light path was used. Image analysis was done with ImageJ software. Hoescht 33342 staining (ThermoFisher) was used to label the nucleus according to manufacturer’s instructions.

### RNA visualization

RNA in live cells was visualized using Pepper RNA aptamer as previously described ^31^. Eight times Pepper repeats were cloned downstream of the T7 promoter and its expression was driven with light starting 24h after transfection. After 30h of illumination, cells were stained using 20 μM of HBC620 supplemented with 5 mM of MgSO_4_. Cells were incubated for 30min at 37 °C with 5% CO_2_ and then transferred to the microscope for imaging.

### Viability assay

Cells were stained using Calcein AM (Sigma-Aldrich) dye after 24h of blue light illumination at different intensities. Cells were prepared for FlowCytometry analysis as described previously. 30 min before measurements Calcein AM was added to the cells at 1 μM final concentration. Samples were incubated on ice until measurement.

### Statistics

Each experiment was repeated with at least three independent biological replicates from which mean and standard deviation are calculated.

## RESULTS

### Characterizing mOptoT7 in mammalian cells

To build a light-inducible system that can function orthogonally to the transcription machinery of mammalians cells, we implemented an optogenetic tool based on a split T7 RNA polymerase fused to the Vivid (VVD) derived Magnet photodimerization system (Opto-T7) previously described in bacteria ^26^. In the presence of blue light, the Magnets dimerize and recognize each other due to electrostatic interactions, ^27^ which leads to the reconstitution of the full, active protein. We tested different versions of the split T7 polymerase and found that these did not function better than the previously published split T7 in terms of fold change (Supplementary Fig. 1a). We, therefore, proceeded with the published split site at the amino acid position 563 and created two separate vectors containing T7(1-563) fused to nMag and T7(564-883) fused to pMag under the control of EF1a promoter, which we call mOptoT7. We built a reporter construct containing a full T7 promoter ^32^ to trigger the expression of mRuby3 and assess the functionality of our optogenetic tool. Due to the viral nature of the T7 polymerase, RNAs produced by the polymerase lack a 5’cap for translation initiation. To allow for initiation of translation, an IRES2 (Internal Ribosome Entry Site type 2) sequence was added between the promoter and the gene of interest (Fig. 1b). We started with the characterization of the system in HEK293T cells by measuring the level of mRuby3 after 24h of constant blue light illumination. We used a previously published^29^ Light Plate Apparatus (LPA), that we optimized for our illumination experiments in mammalian cell culture conditions (see Materials and Methods section and Supplementary Fig. 2). Light activation led to an 8-fold increase in fluorescence compared to the dark control (Fig. 1c). Cell viability was not affected by the light conditions during the experiment as shown by Calcein AM assay. (Supplementary Fig. 3).

We next investigated the response of mOptoT7 to different light intensities. Reporter fluorescence was measured in the whole population after 24h of light illumination using flow cytometry. As was previously observed in the bacterial T7 RNAP ^26^, the cells showed a graded response to light (Fig. 1d).

Compared to other available blue light systems ^33,34^, mOptoT7 shows a higher sensitivity to light, making it an ideal system to use when low light is required for saturating gene expression. To visualize single cell gene expression variability in response to light, we imaged HEK293T cells after 24h of constant light illumination (Fig. 1e). Compared to the dark control, we saw an increased mRuby3 fluorescence, which also shows a larger heterogeneity. This heterogeneity is likely due to transient transfection effects ^35^, but a different degradation rate of the mOptoT7 and/or the reporter between cells cannot be excluded ^36,37^. mRuby3+ population showed a relatively wide distribution, with some cells expressing a very high level of mRuby3 (Fig. 1f). Integration of mOptoT7 in the genome using piggyBac transposase did not result in any activation (data not shown). This is probably due to the inhibition of mOptoT7 initiation and elongation as described previously ^23,38^.

### Optimization of the mammalian OptoT7 system

An important advantage of optogenetic systems is the ability to tune gene expression through different light inputs. To test if we can control the levels of mRuby3 using mOptoT7, we performed a screen with different light programs over 24h illumination period; we also performed a screen with different duration of the light pulse applied, always using a saturating light intensity (Fig. 2a and 2b). Measurements were taken using a flow cytometer. As expected, we observed an increase in the activation of fluorescence that is proportional to the increase of light duration in the cycle, with maximal expression reached under constant light (Fig. 2a). We also observed a change in mRuby3 expression when different duration of light is applied. In particular, the system starts showing light activation with a 3h light pulse and changes its response proportionally to the duration of pulses applied (Fig. 2b).

**Figure 2.**
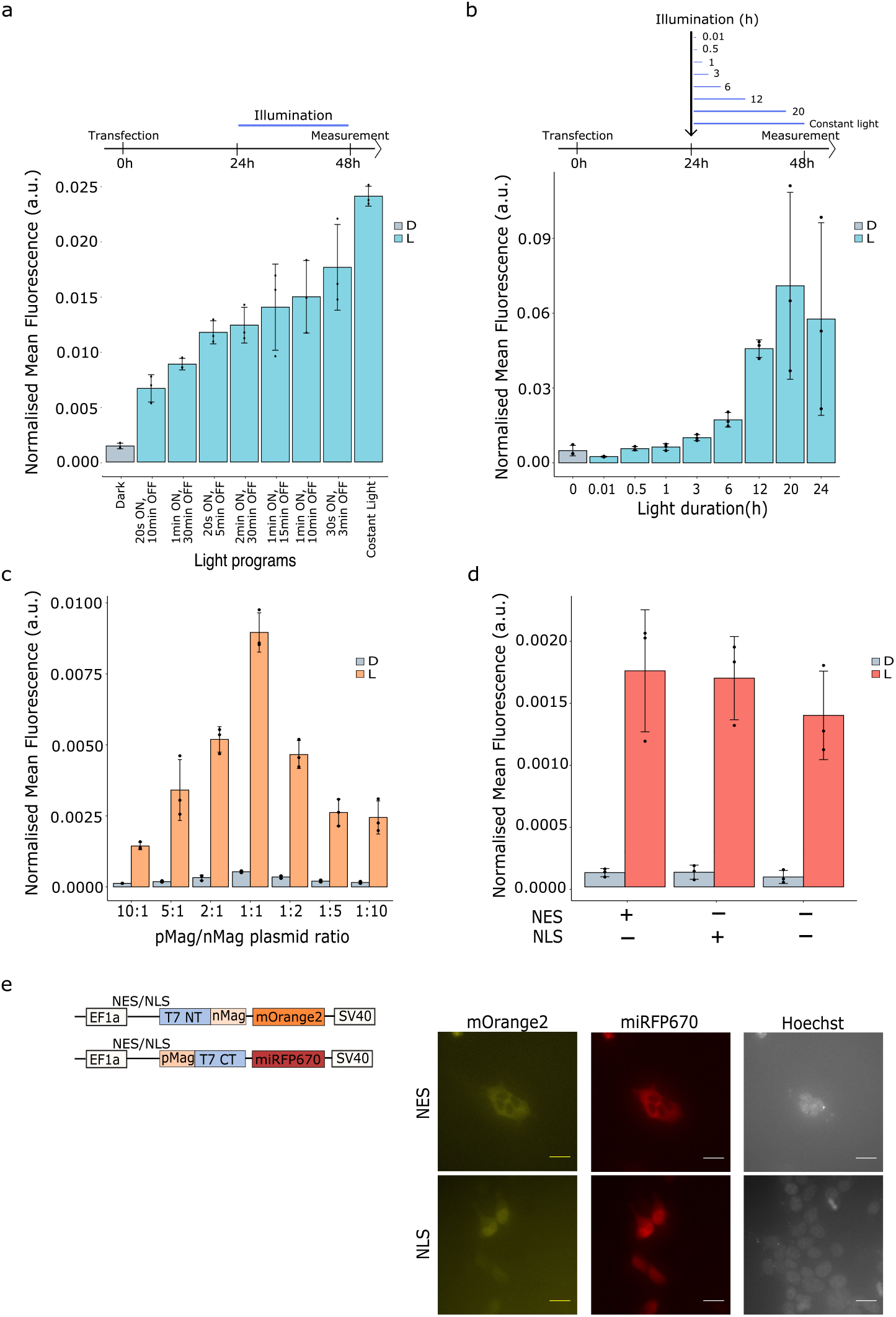
Optimisation of mOpto-T7RNAP in HEK293T cells. (a) Screening through different light conditions using several programs in which the duty cycle duration was changed. Illumination was done for 24h. 2:1 ratio of nMag:pMag was used. Cells were measured at the flowcytometer 24h from illumination. Light was used in saturation regime (200uW/cm2). (b) Screening with different light durations. Cells were measured at the flowcytometer after 48h from transfection. Light was used in saturation regime (200uW/cm2). (c) Screening through different magnets ratios. Displayed is the reporter activation after 24h of constant light illumination under saturating conditions. (d) Characterisation of mOptoT7 with NES (nuclear export sequence) and NLS (nuclear localization sequnce). Reporter expression was measured after 24 of constant light illumination and shows that mOptoT7 can efficienrtly transcribe bobth in the nucleus and in the cytoplasm. (e) Left: Schematics of the constructs used in this experiment. Right: Fluorescence images of mOptoT7 subunits with NES and NLS fused to mOrange2 and miRFP670. Images were taken after 24h from illumination. Hoechst 33342 was used to labbbel the nucleus. Scale bar, 20 micron. L=light, D=dark. Flow cytometry data are normalized to the constitutively expressed mCitrine.

In addition to using different light inputs durations, we sought to exploit the ability to change the ratio of the mOptoT7 fusion proteins during transient transfection experiments (Fig 2c). We observed a variation in both expression level and fold change with changing pMag-/nMag-fusion ratios. In particular, the highest expression level was obtained using 1:1 ratio of the magnets and the highest fold change compared to dark control with an excess of pMag-fusion (5:1). This last finding is consistent with investigations about OptoT7 in bacteria in which an increased expression of the pMag-fusion shows the highest light-induced fold change^26^. Ratiometric control over pMag and nMag levels can thus be used for further fine tuning the mOptoT7 system.

Given the viral origin of the T7 polymerase, mOptoT7 is orthogonal to the cellular machinery for transcription. To execute its function, mOptoT7 only requires nucleotides and Mg(2+) ions, components that can be found both in the nucleus and in the cytoplasm ^22^. We hypothesized that mOptoT7 will be able to transcribe RNA very efficiently outside of the nucleus, allowing a complete separation of transcription activities from the endogenous cellular transcription, and at the same time concentrating mRNA directly in the cytoplasm. To test if mOptoT7 can indeed function within and outside the nucleus, we introduced either a strong nuclear localization sequence (NLS) or a strong nuclear export sequence (NES) before both subunits of the mOptoT7 and compared the activities of these constructs with the original ones without NLS/NES by the measuring mRuby3 fluorescence after 24h of blue light illumination in saturating conditions (Fig. 2d). We observed that fluorescent protein expression level did not significantly change between the two variants, supporting the conclusion that mOptoT7 can transcribe RNA directly and efficiently in the cytoplasm. To confirm the localization of the mOptoT7 in the different cellular compartments, we next fused a fluorescent protein (either mOrange2 or miRFP670) to each of the mOptoT7 subunits, and imaged the fluorescence at the microscope 48h after transfection (Fig. 2e). Supporting our previous results, we observed a strong nuclear or cytoplasmic localization for the variants with NLS and NES sequences respectively, while the mOptoT7 without any localization sequence localized in the whole cell (Supplementary Fig. 4). We did not observe a change in localization after shining constant blue light for 24h nor did we observe significant changes when the fluorescent proteins were switched between the two subunits for the conditions tested (Supplementary Fig. 4).

### Fine-tuning of mOptoT7

As previously mentioned, the T7 polymerase generates RNA transcripts that lack 5’ and 3’UTR modifications. These modifications are essential in controlling translation initiation and increasing RNA stability ^39,40^. Without them, RNA transcripts won’t be translated and RNA half-life will be short. Thus, to further fine-tune OptoT7-mediated reporter expression, we created new variants of the mOptoT7 by changing 5’ and 3’ UTR sequences independently (Fig. 3a).

**Figure 3.**
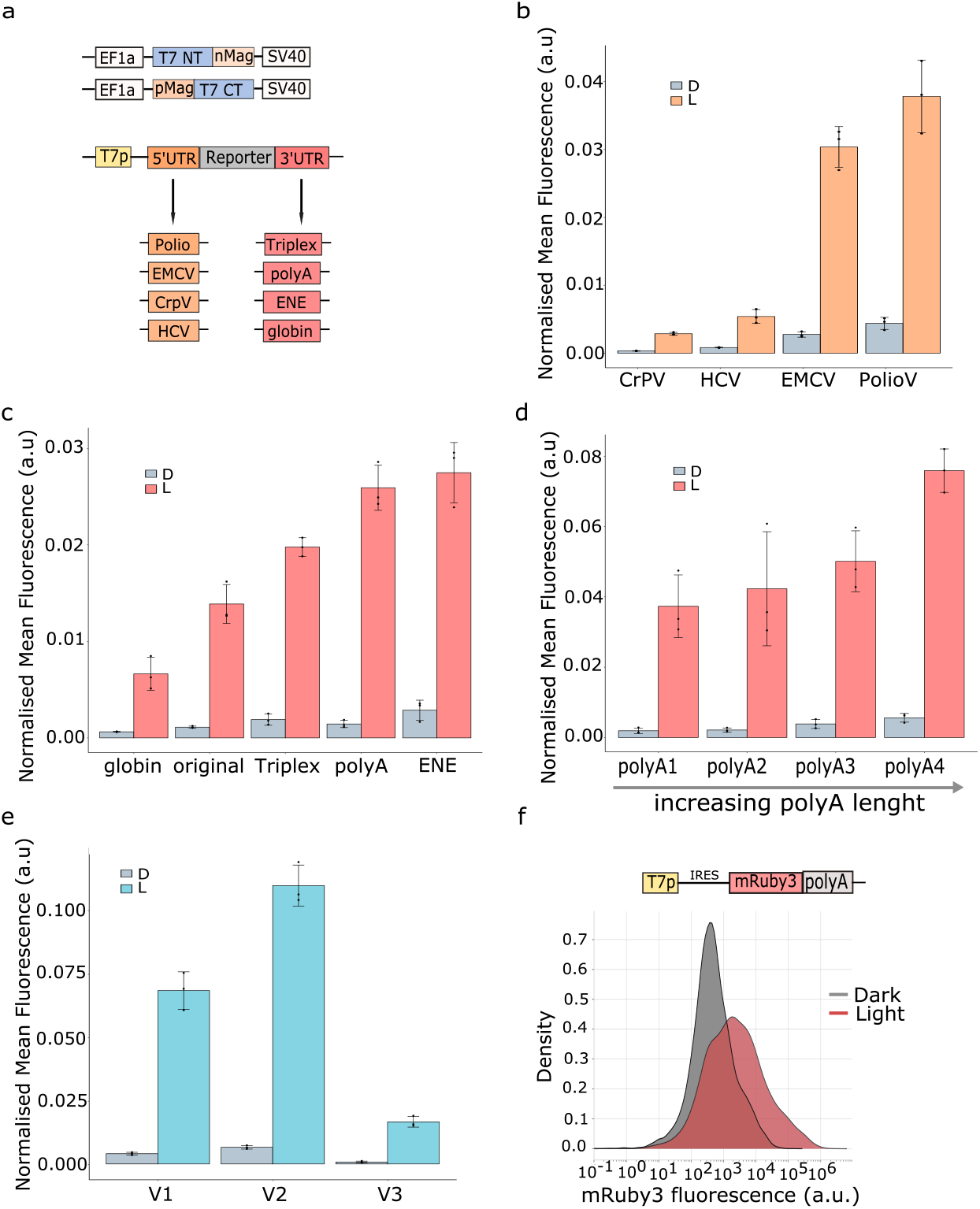
mOptoT7 fine tuning: 3’ and 5’ UTR modifications. (a) Experimental design. OptoT7 is transfected in HEK293T cells together with reporters containing one or more 5’ (shown in orange) and 3’ (shown in red) UTR modifications. (b) Testing of 5’UTR modifications shows different expression levels for different IRES sequences. CrPV= Cricket paralysis virus; HCV= Hepatitis C virus; EMCV= Encephalomyiocarditis virus; PolioV= Poliovirus. (c) Testing of 3’UTR modifications, globin= 3’UTR from human globin gene; original= no 3’UTR modification; Triplex= RNA triple-helical structure; polyA= synthetic poly(A) stretch; ENE= Element for nuclear expression. (d) Characterization of polyA tails with different lengths shows an increase in reporter expression proportional to the lengh of AAAs. (e) New OptoT7 versions. V1= Original design; V2= codon optimized version; V3= shorter 5’UTR sequence. V2 shows the highest expression level while V3 shows tight light response in saturating conditions. D=dark, L=light. Data are normalized to the constitutively expressed mCitrine. For all experiments, measurements were taken 24h after illumination in saturating conditions (200uW/cm2). f) Kernel density estimation plot of mRuby3 expression with the optimized version of mOptoT7. HEK293T cells were transfected with mRuby3-polyA reporter construct and visualized after 24h of constant blue light in saturating conditions 200 μW/cm2.

Inspired by the different strategies that viruses have evolved to initiate translation and stabilize RNA in mammalian cells, we designed new reporter constructs containing IRES sequences from different type of viruses: (i) PolioV-IRES (IRES1) (ii) EMCV-IRES (IRES2) (iii) CrpV-IRES (IRES3/4) and (iv) HCV-IRES (IRES4), upstream of mRuby3 (Fig. 3a) and measured fluorescence after 24h of light illumination (Fig. 3b). These sequences are categorized in four different types according to RNA structure and eIFs (Eukaryotic initiation factors) recruitment, with type 1 being the most complex and recruiting most factors and type 4 having a simple structure and binding directly to ribosomal subunits for translation initiation ^41,42^. We hypothesized that by recruiting different eIFs, IRES sequences can be used to generate different expression levels of mOptoT7 reporter construct. We indeed observed that these structures can titrate different expression levels of mRuby3. Maximum expression was obtained using PolioV-IRES, while the lowest expression was obtained using CrPV-IRES. In particular, due to their simplistic structure and ability to initiate translation with only few translation elongation factors (eIFs), CrPV- and HCV-IRES could be of interest for applications that do not require the use of all cellular resources for translation.

Next, we focused on 3’UTR modifications. In eukaryotic cells, endogenous RNA is made bearing 3’UTR modifications in the form of a poly(A) stretch. This repetition of (A)s is correlated with RNA stability, and transcripts that lack poly(A) tails are known to be short lived ^43^. With the aim of increasing RNA stability and thus, mRuby3 expression in our mOptoT7 reporter, we created constructs that contain different stabilizing sequences at the 3’ region of mRuby3: (i) A triple RNA helix (triplex) that, as described previously ^44^, is used to stabilize noncoding RNAs; (ii) a long stretch of synthetic poly(A)s to mimic the natural occurring adenylation process; (iii) an Element for nuclear expression (ENE) that, as described previously ^45^, is used to stabilize poly(A) transcripts by sequestrating them in triple helix structures; or (iv) the 3’UTR of the globin gene, which is known to be rather stable ^46^ (Fig. 3a). We observed that we could indeed tune reporter expression levels in a wide range after 24h of light illumination by changing the 3’UTR sequence of the mRuby3 reporter, with maximum expression obtained using the synthetic poly(A) stretch, and the lowest expression using the 3’UTR of the globin gene (Fig. 3c). Interestingly, the addition of a poly(A) tail not only increased mRuby3 expression, but also increased the total population that was responsive to light (Fig. 4b, Supplementary Fig. 5). Given the high expression level obtained with the synthetic poly(A) tail, we sought to investigate whether the length of the poly(A) tail corresponds to the reporter stability and could thus be used to further increase mRuby3 expression after 24h of light illumination. As expected, we observed an increase in fluorescence signal with increasing poly(A) length (Fig. 3d).

**Figure 4.**
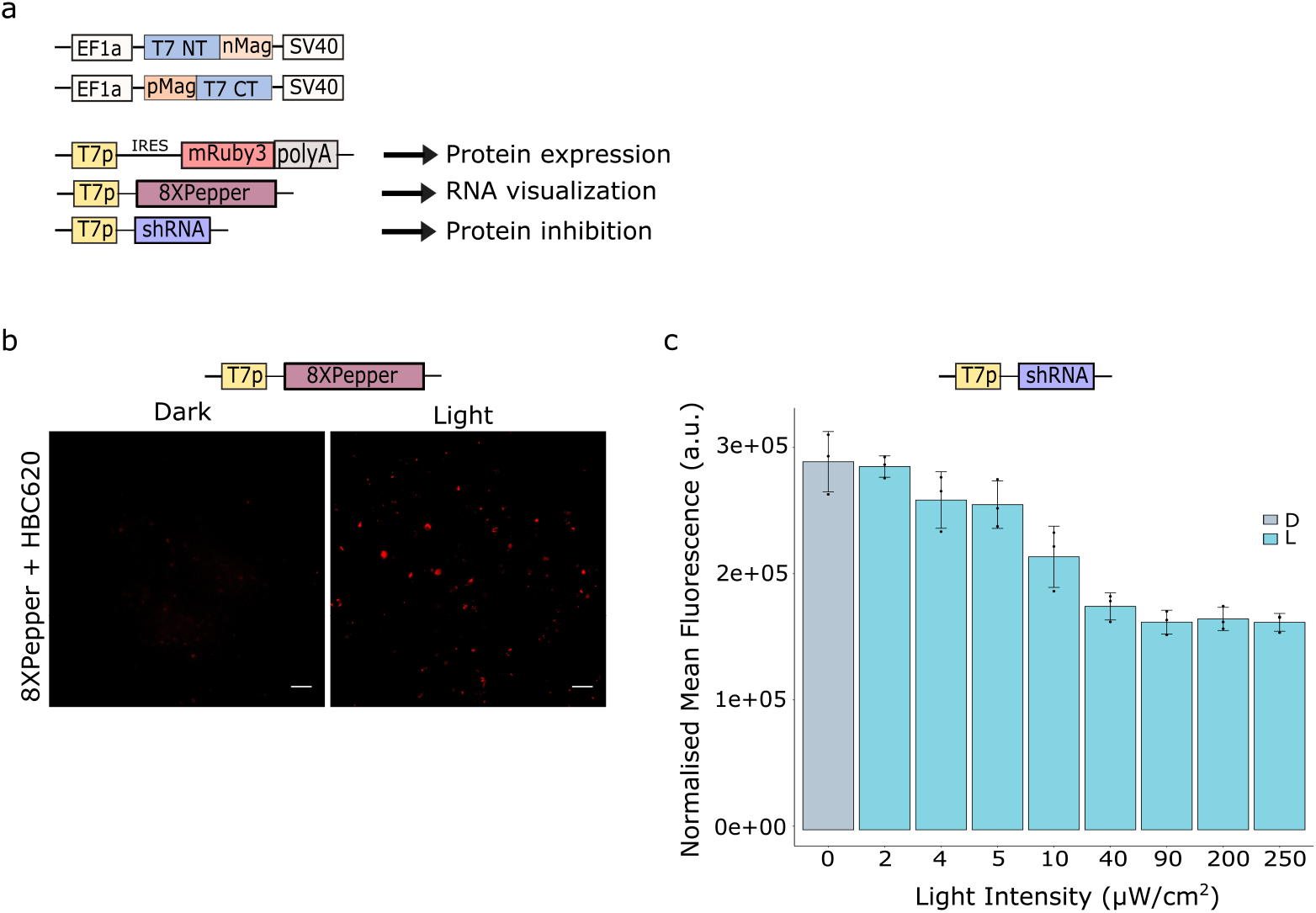
Protein expression/inhbition and RNA visulazion with mOptoT7. (a) Design of the mOptoT7 vectors for gene expression, inhibition and RNA visualization (b) Testing of the mOptoT7 plasmid with a fluorescent RNA aptamer (8XPepper) as the output. Images were taken after 30h of constant blue light illumination. Scale bar, 50 microns. (c) Flowcytometry results of the mOptoT7 vector with shRNA targeting mCitrine as the output. Cells were measured after 24h of constant blue illumination. Data are normalized to the constitutively expressed mCitrine.

Finally, we aimed to create versions of the mOptoT7 that not only had higher light-induced expression levels but also showed reduced leakiness, as an ideal optogenetic system should satisfy both of these criteria. Thus, we first created codon optimized versions of the magnets (V2) that, when combined with the reporter containing the synthetic poly(A) tail gave around 20-fold change and low background activity (Fig. 3e) compared to the non-codon optimized version (V1). Induction of mRuby3 reporter fluorescence with our optimized mOptoT7 version (V2) showed the highest fluorescent protein expression as well as the highest percentage of cells responding to the light input (Fig. 3f). Almost 50% of the population activates mRuby3 production after illumination compared to only 15% using the construct before optimization (Supplementary Fig. 5). We then created another version (V3) by modifying the length of the 5’UTR region of the optimized mOptoT7 polymerase fusion proteins. We hypothesized that the length and composition of 5’UTR sequence will affect transcription rate and RNA stability. Indeed, we found that by decreasing the number of nucleotides between the constitutive EF1a promoter and the start of each subunit of the mOptoT7, we were able to change its transcription and therefore protein availability, to create a very tight gene expression system. When light is applied, this version shows 10-fold change with saturating light and no measurable background activity. This last system is to be preferred in applications for which tight control in the dark is essential while high expression is not required.

### mOptoT7 can be used to visualize RNA and inhibit gene expression

We next assessed the ability of mOptoT7 to generate two more outputs apart from gene expression for a wider range of applications: (i) RNA production for visualization, and (ii) inhibition of gene expression (Fig. 4a). Both RNA visualization and gene expression inhibition were previously shown using T7 polymerase in mammalian cells ^19^, but without the opportunity for dynamic control that is enabled by optogenetics. For our experiments, we created one vector containing both subunits of the mOptoT7 so that we could easily use the 1:1 ratio of the mOptoT7 fusion proteins that showed the highest expression level.

The ability of mOptoT7 to produce RNA species without 5’ and 3’ UTR can be exploited to generate hairpin repeats that can be visualized after binding with fluorophores or used in downstream applications. Thus, we next used a Pepper RNA aptamer (8 repeats) under the control of the T7 promoter to visualize RNA production using the mCherry fluorophore-like synthetic dye HBC620 ^31^. RNA production is detected after 24h of constant light illumination, while the dark control shows no signal (Fig. 4b). Non-transfected cells stained using HBC620 show no expression (Supplementary Fig. 6a). Cells expressing 8xPepper constitutively in the presence or absence of light showed a uniform distribution of RNA within the cell population, and mainly in the cytoplasm (Supplementary Fig. 6b). Interestingly, cells exhibited different patterns of RNA expression, with some cells expressing many RNA molecules homogeneously diffused and others expressing RNA in the form of “dots”. This difference in expression is likely due to different plasmid uptake during transient transfection experiments, however, for some of the bigger dots observed, a stress mechanism that could lead to the formation of RNA granules cannot be excluded. These stress granules were described previously as aggregates that form after stress-induced mechanisms (for example viral infection) and inhibit translation ^47,48^. Potentially, disruption of these granules could further increase mOptoT7 reporter expression level.

Finally, we wanted to investigate if we can also use mOptoT7 to inhibit gene expression. Therefore, we made another vector containing an shRNA hairpin ^49^ against mCitrine under the control of the T7 promoter. We transfected cells with this construct together with the mOptoT7 and a constitutive mCitrine plasmid and measured the expression of mCitrine in response to 24h light exposure at different intensities (Fig. 4c). We observed that the fluorescent signal decreased with increasing light intensity, reaching almost 50% of the mCitrine fluorescence levels in the dark. These results show that mOptoT7 can also be used to inhibit gene expression through a polymerase that is orthogonal to the cellular one, highlighting the potential for the use of mOptoT7 for research questions on gene function.

### mOptoT7 shows reduced burden on the host cell

Finally, we wanted to assess the ability of mOptoT7 to avoid gene expression burden that is commonly exerted by other optogenetic tools that rely on the recruitment of the cellular polymerase. Given the orthogonality of mOptoT7 in generating RNA transcripts (Fig. 2d), we hypothesized that mOptoT7 will use less overall cellular resources (e.g. polymerase subunits, eIFs), thereby imposing less burden to the cell.

To test this, we used mOptoT7 together with the reporter bearing the IRES2 sequence and the longest poly(A) tail to have the highest expression level of mRuby3. To measure the effect of increasing mRuby3 expression level in response to light, we used a “sensor gene” called capacity monitor as previously described ^8^. In our case, the capacity monitor consisted of mCitrine fluorescent protein under the control of EF1a promoter (Fig. 5a). As comparison, we used another VVD-based optogenetic tool, called GAVPO ^50^. Given that GAVPO has a stronger reporter expression with light compared to mOptoT7, we replaced the p65 activation domain with the weaker activation domain VP16 ^51^ (Fig. 5b). In response to increasing light intensity, we observed an increase in reporter expression for both mOpotoT7 and GAV-VP16, with the latter having weaker mRuby3 expression level (Fig. 5b). Interestingly, when looking at the capacity monitor for both systems, we clearly saw a sharper decrease in mCitrine expression level with increasing light intensity when using GAV-VP16, despite having a weaker expression level for the reporter (Fig. 5c, right) compared to the mOptoT7 reporter expression (Fig. 5C, left). This supports our hypothesis that mOptoT7 can be used to minimize gene expression burden. To exclude the possibility that blue light causes this effect, we shined an increasing amount of light on cells transfected only with the capacity monitor and measured their expression level after 24h from illumination. No decrease in mCitrine levels was observed at the applied light intensities (Supplementary Fig. 7).

**Figure 5.**
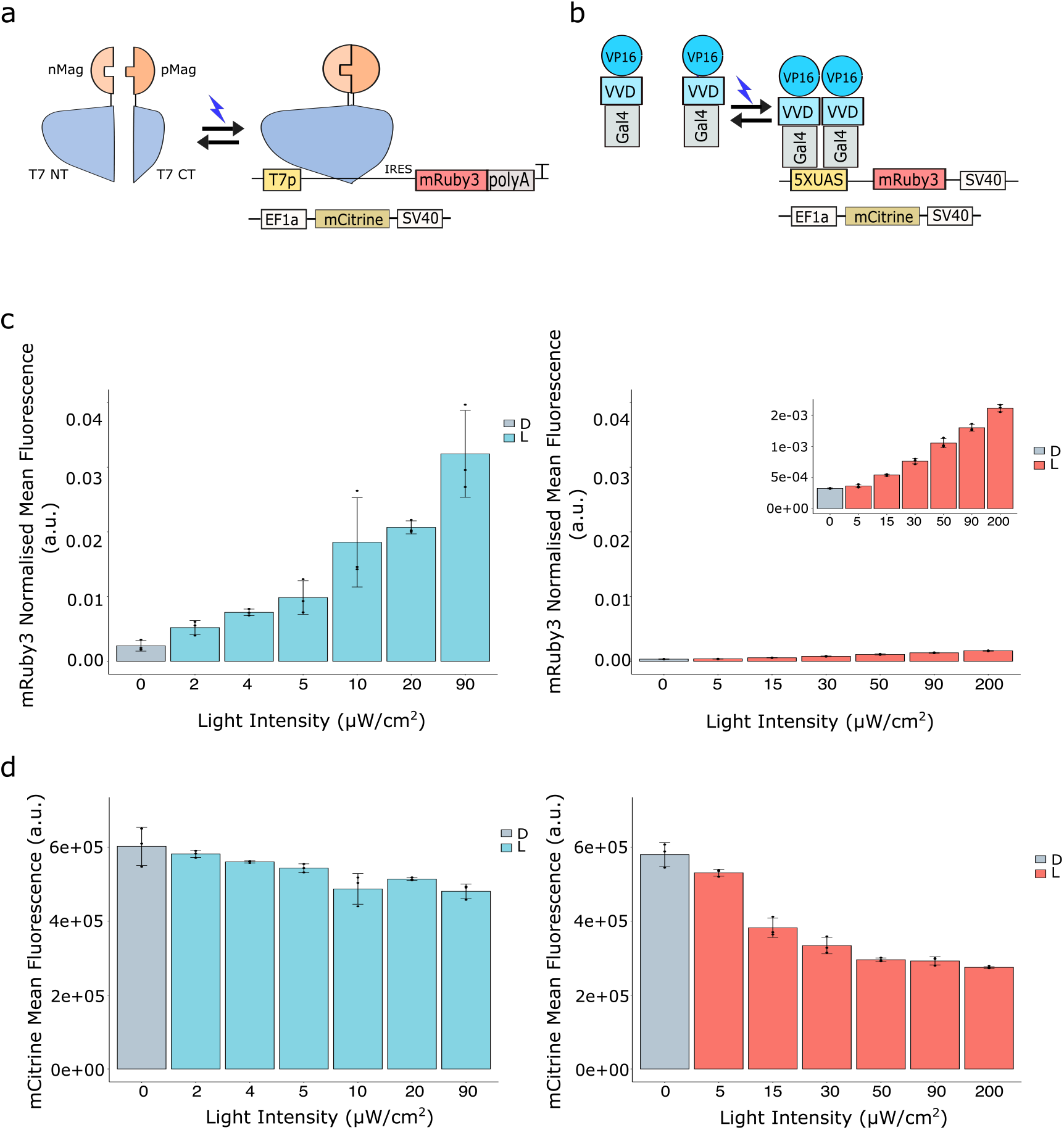
mOptoT7 mitigates burden in HEK293T cells. (a) and (b) Schematics of mOptoT7 and GAV-VP16 used in this experiment. mRuby3 fluorescent protein was used as reporter color. mCitrine under the control of EFla promoter was used as constitutive color (capacirty monitor).(c) Dose response of mOptoT7 (left panel) and GAV-VP16 (right panel) reporter to increasing light intensity. (d) Capacity monitor’s response to increasing light intensity for mOptoT7 (left) and GAV-VP16 (right) shows less reduction in the expression of mOptoT7 capacity monitor compared to GAV-VP16. D=dark, L=light. Data are normalized to the constitutively expressed mCitrine. Measurements are taken after 24h of constant blue light illumination.

## DISCUSSION

In this study we created mOptoT7, a novel optogenetic tool in HEK293T cells based on a split T7 RNA polymerase coupled with the light-responsive magnet dimers derived from VVD ^26^. This is the first time that an optogenetic system orthogonal to the cellular transcriptional machinery is applied in mammalian cells. mOptoT7 can activate downstream gene expression upon light exposure with almost 20-fold change over the dark control and relatively low leakiness (Fig. 3e). By changing the 5’ and 3’ UTR ends of the RNA species generated, we were able to fine-tune mOptoT7 reporter expression creating a wide range of induction responses. The addition of poly(A) tails at the 3’UTR of the reporter construct not only increased expression level, but also the number of cells that were activated after illumination (Fig. 4b).

While we observed heterogeneity in reporter gene expression upon light activation of mOptoT7, this is not unlike other optogenetic tools, especially during transient transfection experiments ^34,35,52^. On a more general level, little is known about the way optogenetic systems affect cellular functions or how they are affected by cellular state and metabolic stress, complicating their robust use in different cell lines and conditions ^53,54^. Future work should focus on solving some of these problems.

We showed that mOptoT7 can efficiently transcribe RNA both in the cytoplasm and in the nucleus. This property is particularly relevant when compartmentalization of RNA is desired. Battich et al. showed that nuclear retention of RNA can filter out noise and explain transcriptional bursts in mammalian cells ^55^. However, until now, no system is available to investigate and potentially control how the noise distribution is affected by RNA production in different compartments, which could also have an application in building synthetic cells^56^.

The capability of mOptoT7 to transcribe RNA in the cytoplasm is connected to its orthogonality to the cellular machinery. This property is especially interesting if applied to study resource allocation and gene expression burden. Recently, several studies have shown how burden in mammalian cells is generated through the sharing of cellular resources both at the transcriptional and translational level ^8,9^. We showed that using mOptoT7 to orthogonally generate RNA species (in an inducible manner) not only helps in mitigating burden, but can also help in gaining a better understanding of how transcriptional resources affect burden as a whole. Contribution of IRES sequences that are included in the mOptoT7 reporter are not considered in this study and require further investigation.

Besides orthogonality, another interesting feature of mOptoT7 is its ability to generate transcripts that lack endogenous RNA modifications. In this study we visualize the localization of these transcripts using Pepper RNA aptamers. Interestingly, we see dot-like structures located mostly around the nucleus. These aggregates could be virus-induced RNA granules that are formed in response to viral infection ^47,48^; however this remains a theory at this point. As such, mOptoT7 could be employed to study viral RNA recognition and degradation in mammalian cells.

Studies in bacteria have shown how the T7 RNA polymerase can be used to create more robust and stable genetic circuits ^16,57^. The ability of mOptoT7 to be induced with light, as well as the diversity of available T7 variants can be used as powerful tool to create synthetic networks with different feedback and feedforward properties. These genetic circuits can be used for cell-based therapies and clinical applications where stability and reliability play an important role.

To conclude, in this study we implemented, characterized and optimized a novel optogenetic tool in mammalian cells. mOptoT7 has some unique features that are not shared by any other optogenetic tool currently available. The orthogonality to the cellular machinery and the viral RNA origin can be exploited both in synthetic biology and basic science to gain a better understanding of gene expression processes in mammalian cells.

## DATA AVAILABILITY

All relevant data are included are available from the corresponding author. Strains and plasmids used in this study are available from the corresponding author on reasonable request. Main plasmid maps used can be found in the Supplementary material.

## ACKNOWLEDGEMENTS

We thank Peter Buchmann for his help in building the LED setup, Dr. Stephanie Aoki and Dr. Maaike Welling for their feedback on the manuscript. This project has received funding from the European Research Council (ERC).

## AUTHOR INFORMATION

### CONTRIBUTIONS

S.D. conceived the study, performed experiments, analyzed data, and wrote the paper. AB helped conceive the study, co-supervised KP and reviewed the manuscript. K.P. created the original reporter vector and performed initial characterization experiments. M.K. supervised the study, reviewed the manuscript and secured funding.

