## Supplementary Information for "Implementation of a novel optogenetic tool in mammalian cells based on a split T7 RNA polymerase"

1 Department of Biosystems Science and Engineering, ETH Zürich
*Corresponding author

### Content


- Supplementary Fig.1: Test on different T7 split sites
- Supplementary Fig.2: LPA cage design and optimization
- Supplementary Fig.3: Calcein AM assay
- Supplementary Fig.4: mOptoT7 fusion proteins
- Supplementary Fig.5: Example of gating
- Supplementary Fig.6: Pepper RNA aptamer
- Supplementary Fig.7: Light and burden

- Plasmids sequences:
  1. mOptoT7 Version 1 (V1)
  2. mOptoT7 Version 2 (V2)
  3. mOptoT7 Version 3 (V3)
  4. mOptoT7 reporter
  5. mOptoT7 reporter with polyA
  6. mOptoT7 reporter with 8XPepper RNA aptamer
  7. mOptoT7 reporter with shRNA

a

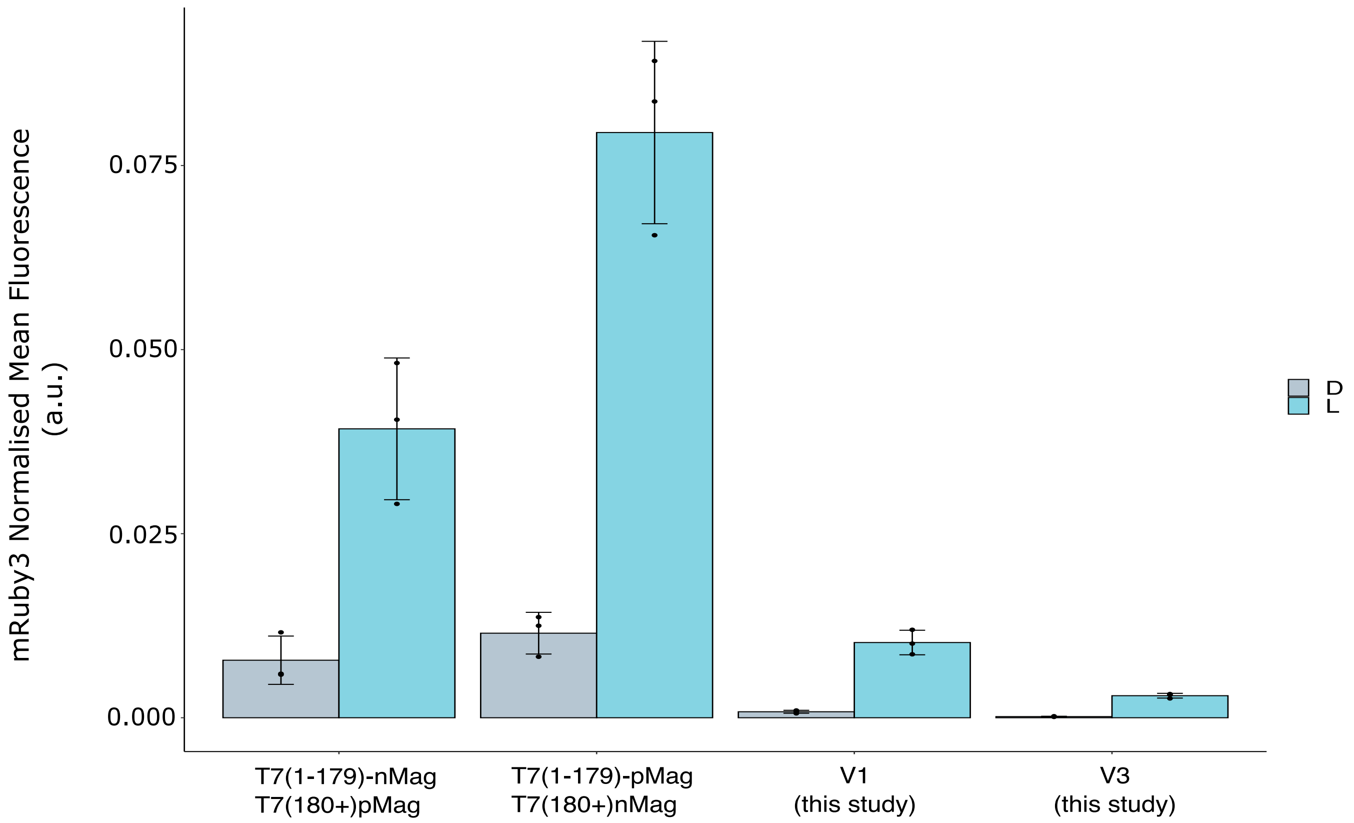


Supplementary Fig.1. mOptoT7 with different split sites. (a) Example of 2 split variants tested in this study. mRuby3 reporter mean fluorescence was measured at the FlowCytometer 24h after illumination. More split sites in the T7 polymerase were tested with negative results (data not shown). T7 polymerase fusion with other blue light optogenetic tools also led to a non-functional tool (data not shown).

a

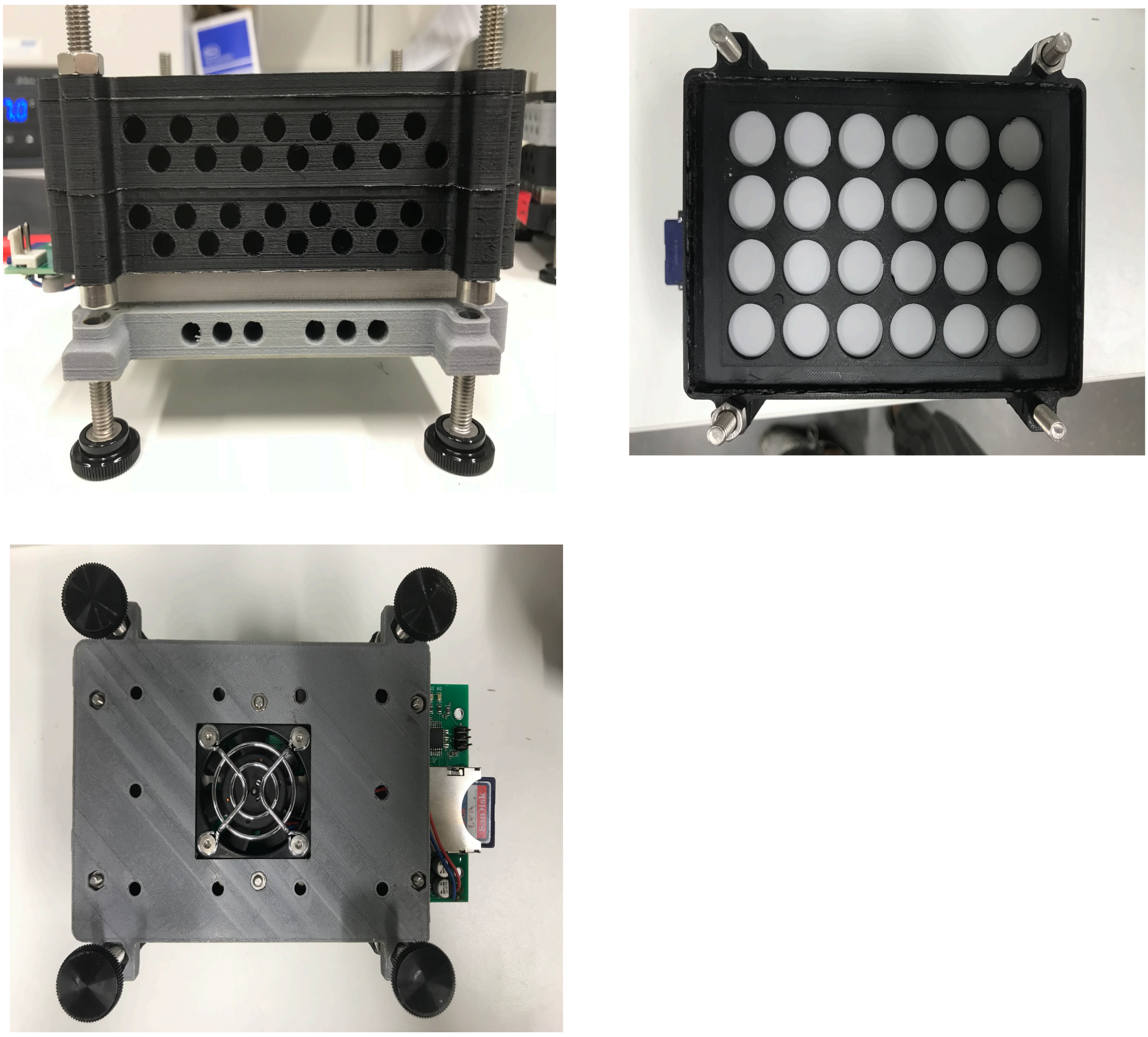

Supplementary Fig.2. Optimization of LPA (light Plate Apparatus). (a) Images of the optimized LPA used in this study. Two black adaptors were used to maximize the distance from the LEDs; a 2mm alluminium heastsink was added between the addaptors and the mounting plate (top left). Filter paper was placed between the two adaptors and between the adaptor and the plate holder to gurantee an homogenous distribution of light (top right). Finally, a ventilator was added at the bottom of the LPA to dissipate the heat from the PCB (bottom left).

a

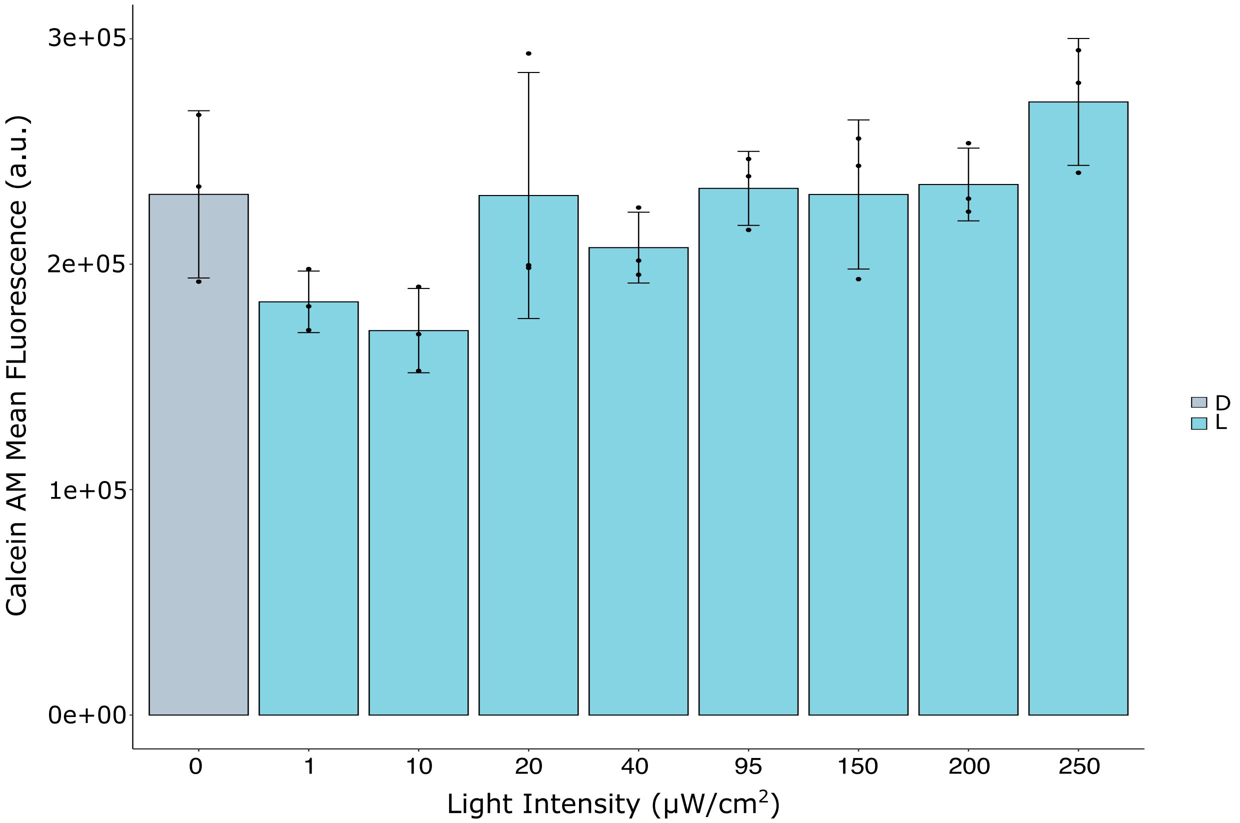


Supplementary Fig.3. Cell viability in response to light. (a) Mean fluorescence of Calcein AM dye after 24h of constant light illumination at different intensities. No toxicity in terms of viable cells was seen between the dark control and the samples exposed to light.

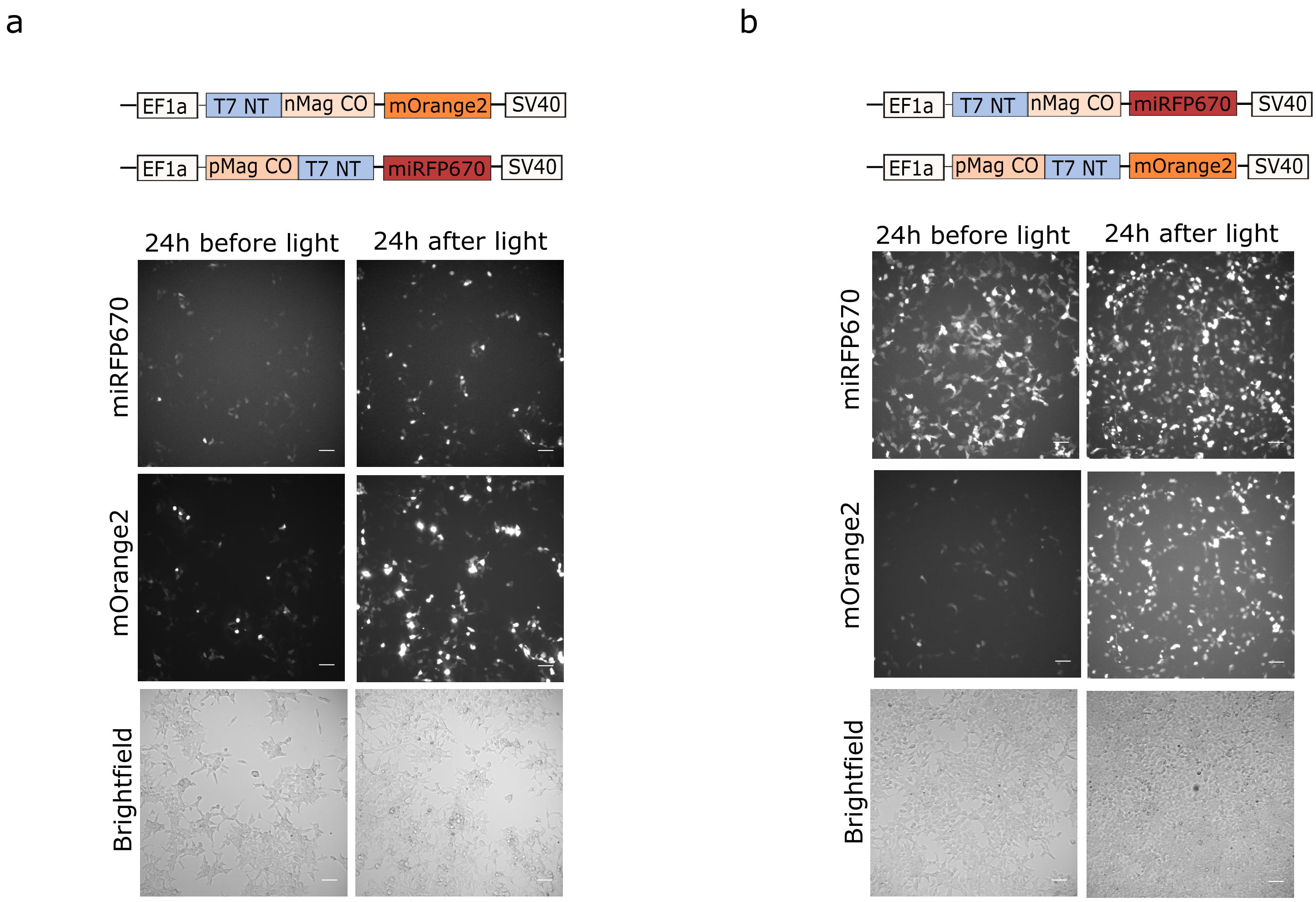


Supplementary Fig4. mOptoT7 fusion proteins. (a-b) Microscopy images of mOptoT7 subunits fused to mOrange2 and miRFP670. No changes in cellular localization was observed after switching fluorescent proteins and after illumination with saturating conditions. CO: Codon Optimised. Illumination started 24h after transfection.

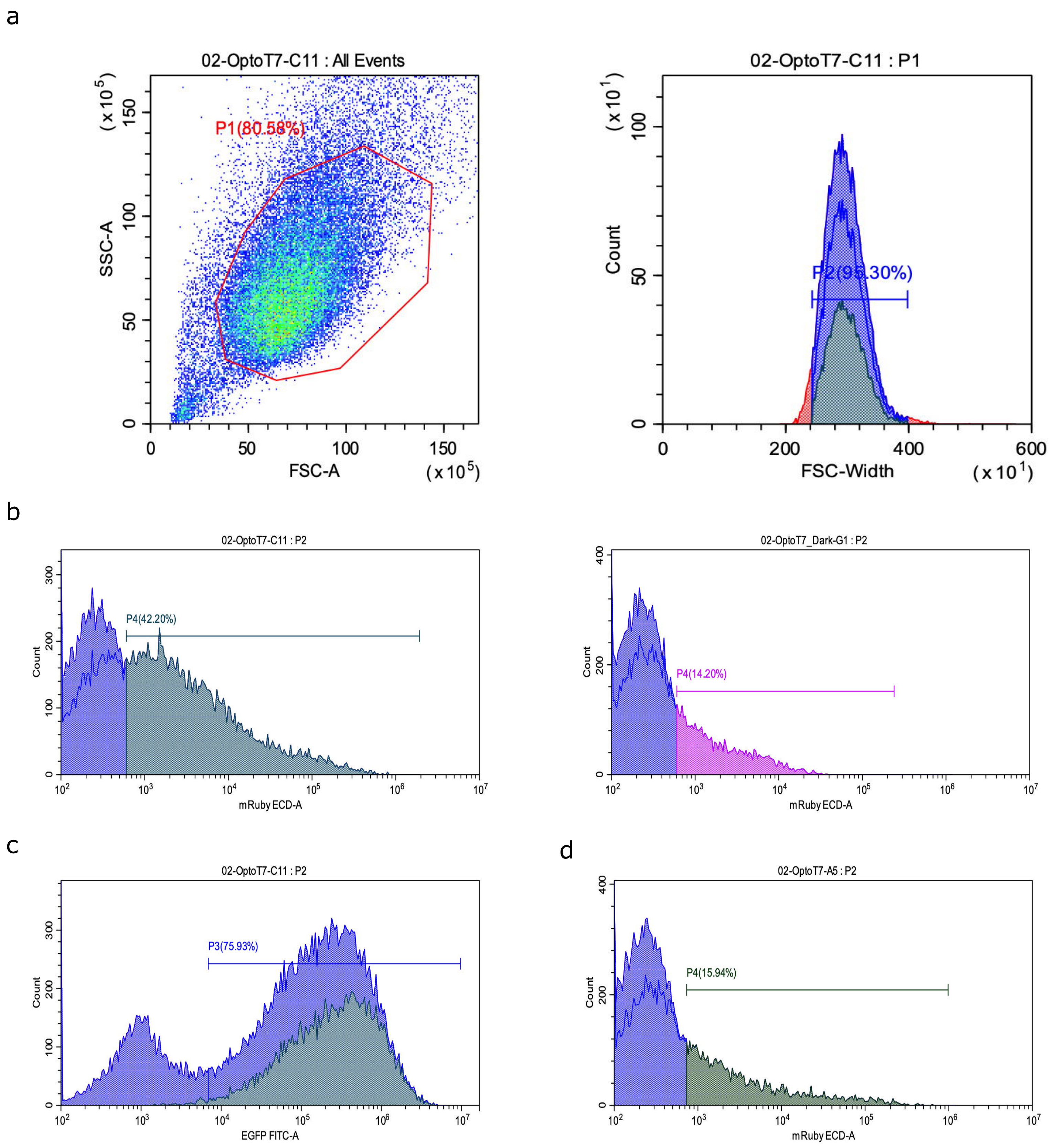


Supplementary Fig5. mOptoT7 gating. (a) Flow Cytometry raw data showing an example of gating for cell population (left) and singlets (right). Cells were transfected with mOptoT7-fusion plasmids and the optimised reporter containing mRuby3 fluorescent protein and polyA tail at the 3'UTR. Measurements were done after 24h from illumination. (b) Histogram of mRuby3 fluorescence reporter shows the percentage of activated cells after light input (left). Dark control shows the number of cells activated in absence of light (right). (c) Histogram of the constitutively expressed mCitrine fluorescence for the same sample. Transfected cells are marked with P3 population. (d) Separate experiment showing mRuby3 fluorescence expression after 24h of light illumination for mOptoT7-fusion proteins and reporter without polyA tail.

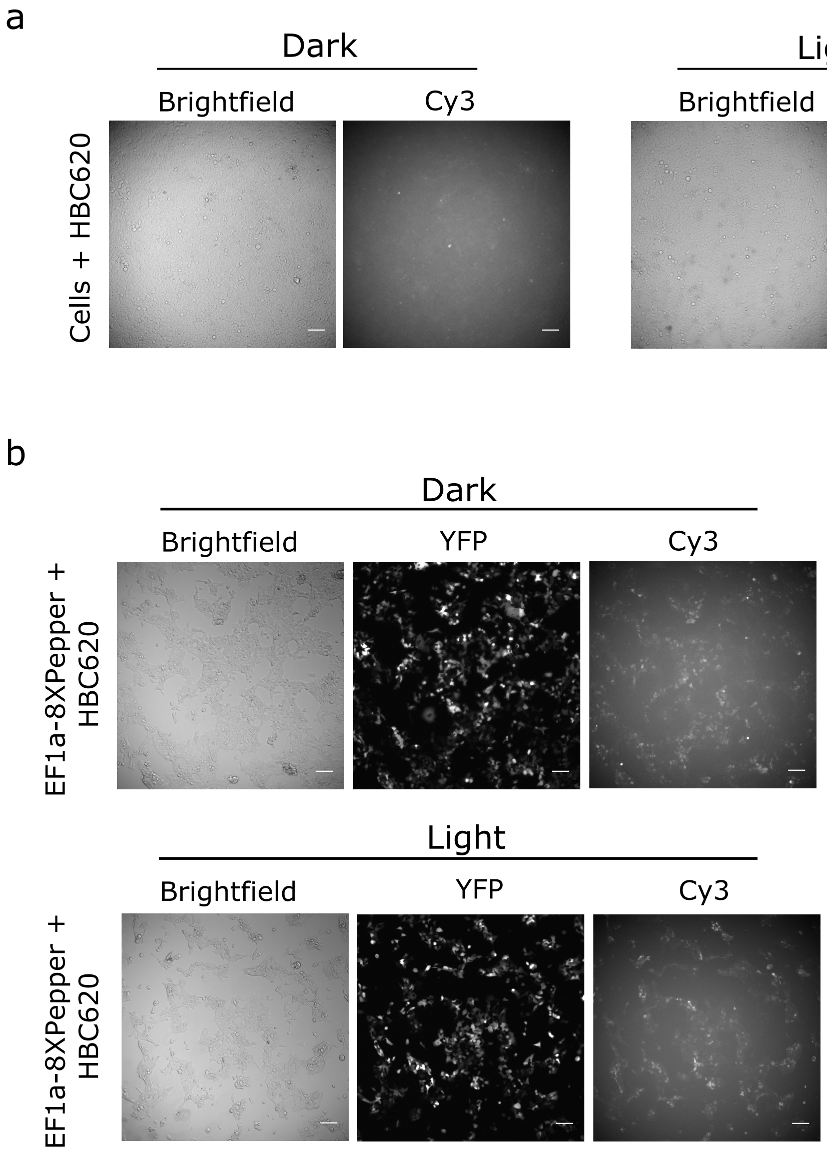


Supplementary Fig6. 8XPepper aptamer for RNA visualization. (a) HEK293T cells stained with HBC620 ligand in absence and presence of blue light. Background fluorescence did not change between conditions. (b) Constitutively expressed 8XPepper RNA aptamer after HBC620 addition in absence and presence of blue light. RNA is sucessufully detected and mainly localises in the cytoplasm. Scale bar, 100 micron.

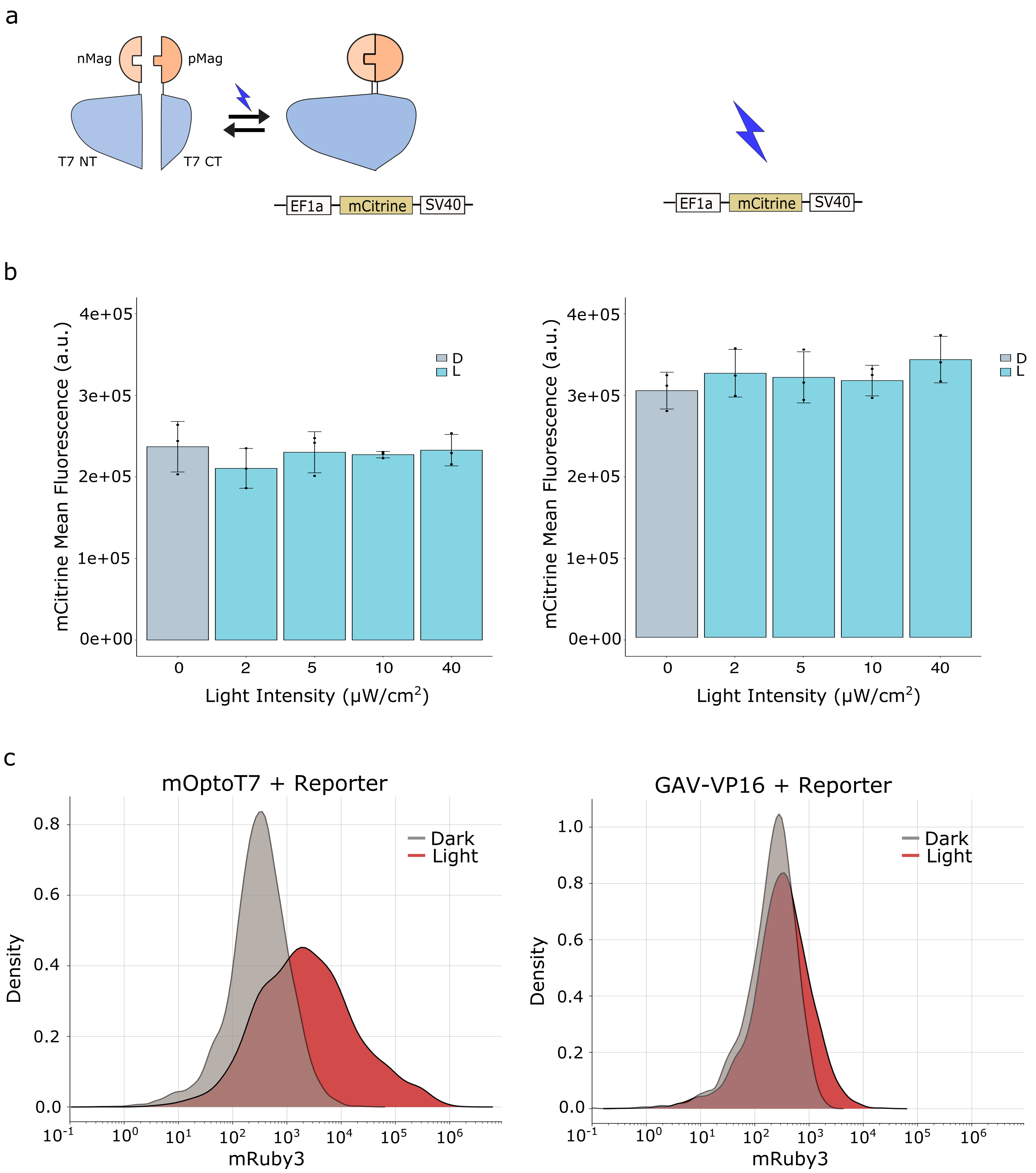


Supplementary Fig.7. mOptoT7 to mitigate burden. (a) Schematics of constructs used in this experiment. mCitrine fluorescence was measured 24h after illumination together with mOptoT7-fusion plasmids (left panel) or alone (right panel). (b) FlowCytometry data shows no effect of increasing light intensity on mCitrine fluorescence when co-transfetcted with mOptoT7-fusion plasmids (left) or alone (right). (c) Kernel density estimation plot showing mRuy3 reporter expression in the dark vs saturating light for mOptoT7 and the engineered weak version of GAVPO (GAV-VP16) used in the study. D=dark, L=light.

PLASMID SEQUENCE

1. mOptoT7 Version 1 (V1)

EF1a promoter
pMag
T7pol (564-883)
SV40polyA

GCTCCGGTGCCCGTCAGTGGGCAGAGCGCACATCGCCCACAGTCCCCGAGAA
GTTGGGGGGAGGGGTCGGCAATTGAACCGGTGCCTAGAGAAGGTGGCGCGG
GGTAAACTGGGAAAGTGATGTCGTGTACTGGCTCCGCCTTTTTCCCGAGGGTG
GGGGAGAACCGTATATAAGTGCAGTAGTCGCCGTGAACGTTCTTTTTCGCAAC
GGGTTTGCCGCCAGAACACAGGTAAGTGCCGTGTGTGGTTCCCGCGGGCCTGG
CCTCTTTACGGGTTATGGCCCTTGCGTGCCTTGAATTACTTCCACGCCCCTGGCT
GCAGTACGTGATTCTTGATCCCGAGCTTCGGGTTGGAAGTGGGTGGGAGAGTT
CGAGGCCTTGCGCTTAAGGAGCCCCTTCGCCTCGTGCTTGAGTTGAGGCCTGG
CCTGGGCGCTGGGGCCGCCGCGTGCGAATCTGGTGGCACCTTCGCGCCTGTCT
CGCTGCTTTCGATAAGTCTCTAGCCATTTAAAATTTTTGATGACCTGCTGCGACG
CTTTTTTTCTGGCAAGATAGTCTTGTAAATGCGGGCCAAGATCTGCACACTGGT
ATTTCGGTTTTTGGGGCCGCGGGCGGCGACGGGGCCCGTGCGTCCCAGCGCA
CATGTTCGGCGAGGCGGGGCCTGCGAGCGCGGCCACCGAGAATCGGACGGG
GGTAGTCTCAAGCTGGCCGGCCTGCTCTGGTGCCTGGCCTCGCGCCGCCGTGT
ATCGCCCCGCCCTGGGCGGCAAGGCTGGCCCGGTCGGCACCAGTTGCGTGAG
CGGAAAGATGGCCGCTTCCCGGCCCTGCTGCAGGGAGCTCAAAATGGAGGAC
GCGGCGCTCGGGAGAGCGGGCGGGTGAGTCACCCACACAAAGGAAAAGGGC
CTTTCCGTCCTCAGCCGTCGCTTCATGTGACTCCACGGAGTACCGGGCGCCGTC
CAGGCACCTCGATTAGTTCTCGAGCTTTTGGAGTACGTCGTCTTTAGGTTGGGG
GGAGGGGTTTTATGCGATGGAGTTTCCCCACACTGAGTGGGTGGAGACTGAAG
TTAGGCCAGCTTGGCACTTGATGTAATTCTCCTTGGAATTTGCCCTTTTTGAGTTT
GGATCTTGGTTCATTCTCAAGCCTCAGACAGTGGTTCAAAGTTTTTTTCTTCCAT
TTCAGGTGTCGTGACATAAACTGCCCTAGAttaatTAACGGCCGCCGCCGCCACC
ATGCACACTCTTTACGCCcctggaggatacgacattatgggatatttgcGGCAGATTAGGA

ACCGCCCAAACCCTCAGGTCGAACTGGGGCCTGTGGACACGTCATGTGCCCTG
ATCCTGTGCGATCTGAAGCAAAAGGACACTCCGATCGTCTACGCCTCGGAAGC
CTTCTTGTATATGACCGGATACAGCAATGCAGAGGTGCTCGGCAGGAACTGCA
GATTCCTGCAGTCCCCCGACGGGATGGTGAAACCAAAGTCGACTCGCAAATAT
GTGGACTCGAACACGATCAACACCATGCGGAAGGCCATCGACCGGAACGCCG
AGGTCCAGGTGGAGGTGGTCAACTTTAAGAAGAACGGCCAGCGGTTCGTGAA
CTTTCTGACCATGATTCCGGTCCGGGATGAAACCGGAGAGTACAGATACTCCAT
GGGATTCCAGTGCGAAACCGAAggcggtTCTggaggtAGTGAAACCGTTCAGGAC
ATCTACGGGATTGTTGCTAAGAAAGTCAACGAGATTCTACAAGCAGACGCAAT
CAATGGGACCGATAACGAAGTAGTTACCGTGACCGATGAGAACACTGGTGAA
ATCTCTGAGAAAGTCAAGCTGGGCACTAAGGCACTGGCTGGTCAATGGCTGGC
TTACGGTGTTACTCGCAGTGTGACTAAGCGTTCAGTCATGACGCTGGCTTACGG
GTCCAAAGAGTTCGGCTTCCGTCAACAAGTGCTGGAAGATACCATTCAGCCAG
CTATTGATTCCGGCAAGGGTCTGATGTTCACTCAGCCGAATCAGGCTGCTGGAT
ACATGGCTAAGCTGATTTGGGAATCTGTGAGCGTGACGGTGGTAGCTGCGGTT
GAAGCAATGAACTGGCTTAAGTCTGCTGCTAAGCTGCTGGCTGCTGAGGTCAA
AGATAAGAAGACTGGAGAGATTCTTCGCAAGCGTTGCGCTGTGCATTGGGTAA
CTCCTGATGGTTTCCCTGTGTGGCAGGAATACAAGAAGCCTATTCAGACGCGCT
TGAACCTGATGTTCCTCGGTCAGTTCCGCTTACAGCCTACCATTAACACCAACA
AAGATAGCGAGATTGATGCACACAAACAGGAGTCTGGTATCGCTCCTAACTTT
GTACACAGCCAAGACGGTAGCCACCTTCGTAAGACTGTAGTGTGGGCACACGA
GAAGTACGGAATCGAATCTTTTGCACTGATTCACGACTCCTTCGGTACCATTCC
GGCTGACGCTGCGAACCTGTTCAAAGCAGTGCGCGAAACTATGGTTGACACAT
ATGAGTCTTGTGATGTACTGGCTGATTTCTACGACCAGTTCGCTGACCAGTTGC
ACGAGTCTCAATTGGACAAAATGCCAGCACTTCCGGCTAAAGGTAACTTGAAC
CTCCGTGACATCTTAGAGTcggacttcgcgtTCGCGTAATCTAGAGGCATCActagtat
gtacaagTAGTAATCTAGAGGGCCCTATTCTATAGTGTCACCTAAATGCTAGAGC
TCGCTGATCAGTCCGTCGACggatccaccggatctagataactgatcataatcagccataccac
atttgtagaggttttacttgctttaaaaaacctcccacacctccccctgaacctgaaacataaaatgaatgc
aattgttgttgttaacttgtttattgcagcttataatggttacaaataaagcaatagcatcacaaatttcacaa
ataaagcatttttttcactgcattctagttgtggtttgtccaaactcatcaatgtatctta

EF1a promoter
nMagHigh1

T7pol (1-563)
SV40polyA

GCTCCGGTGCCCGTCAGTGGGCAGAGCGCACATCGCCCACAGTCCCCGAGAA
GTTGGGGGGAGGGGTCGGCAATTGAACCGGTGCCTAGAGAAGGTGGCGCGG
GGTAAACTGGGAAAGTGATGTCGTGTACTGGCTCCGCCTTTTTCCCGAGGGTG
GGGGAGAACCGTATATAAGTGCAGTAGTCGCCGTGAACGTTCTTTTTCGCAAC
GGGTTTGCCGCCAGAACACAGGTAAGTGCCGTGTGTGGTTCCCGCGGGCCTGG
CCTCTTTACGGGTTATGGCCCTTGCGTGCCTTGAATTACTTCCACGCCCCTGGCT
GCAGTACGTGATTCTTGATCCCGAGCTTCGGGTTGGAAGTGGGTGGGAGAGTT
CGAGGCCTTGCGCTTAAGGAGCCCCTTCGCCTCGTGCTTGAGTTGAGGCCTGG
CCTGGGCGCTGGGGCCGCCGCGTGCGAATCTGGTGGCACCTTCGCGCCTGTCT
CGCTGCTTTCGATAAGTCTCTAGCCATTTAAAATTTTTGATGACCTGCTGCGACG
CTTTTTTTCTGGCAAGATAGTCTTGTAAATGCGGGCCAAGATCTGCACACTGGT
ATTTCGGTTTTTGGGGCCGCGGGCGGCGACGGGGCCCGTGCGTCCCAGCGCA
CATGTTCGGCGAGGCGGGGCCTGCGAGCGCGGCCACCGAGAATCGGACGGG
GGTAGTCTCAAGCTGGCCGGCCTGCTCTGGTGCCTGGCCTCGCGCCGCCGTGT
ATCGCCCCGCCCTGGGCGGCAAGGCTGGCCCGGTCGGCACCAGTTGCGTGAG
CGGAAAGATGGCCGCTTCCCGGCCCTGCTGCAGGGAGCTCAAAATGGAGGAC
GCGGCGCTCGGGAGAGCGGGCGGGTGAGTCACCCACACAAAGGAAAAGGGC
CTTTCCGTCCTCAGCCGTCGCTTCATGTGACTCCACGGAGTACCGGGCGCCGTC
CAGGCACCTCGATTAGTTCTCGAGCTTTTGGAGTACGTCGTCTTTAGGTTGGGG
GGAGGGGTTTTATGCGATGGAGTTTCCCCACACTGAGTGGGTGGAGACTGAAG
TTAGGCCAGCTTGGCACTTGATGTAATTCTCCTTGGAATTTGCCCTTTTTGAGTT
TGGATCTTGGTTCATTCTCAAGCCTCAGACAGTGGTTCAAAGTTTTTTTCTTCCA
TTTCAGGTGTCGTGACATAAACTGCCCTAGAttaattaaCTAGTGCGGCCGCCGCC
GCCACCATGAACACGATTAACATCGCTAAGAACGACTTCTCTGACATCGAACT
GGCTGCTATCCCGTTCAACACTCTGGCTGACCATTACGGTGAGCGTTTAGCTCG
CGAACAGTTGGCCCTTGAGCATGAGTCTTACGAGATGGGTGAAGCACGCTTCC
GCAAGATGTTTGAGCGTCAACTTAAAGCTGGTGAGGTTGCGGATAACGCTGCC
GCCAAGCCTCTCATCACTACCCTACTCCCTAAGATGATTGCACGCATCAACGAC
TGGTTTGAGGAAGTGAAAGCTAAGCGCGGCAAGCGCCCGACAGCCTTCCAGTT
CCTGCAAGAAATCAAGCCGGAAGCCGTAGCGTACATCACCATTAAGACCACTC
TGGCTTGCCTAACCAGTGCTGACAATACAACCGTTCAGGCTGTAGCAAGCGCA

ATCGGTCGGGCCATTGAGGACGAGGCTCGCTTCGGTCGTATCCGTGACCTTGA
AGCTAAGCACTTCAAGAAAAACGTTGAGGAACAACTCAACAAGCGCGTAGGG
CACGTCTACAAGAAAGCATTTATGCAAGTTGTCGAGGCTGACATGCTCTCTAAG
GGTCTACTCGGTGGCGAGGCGTGGTCTTCGTGGCATAAGGAAGACTCTATTCA
TGTAGGAGTACGCTGCATCGAGATGCTCATTGAGTCAACCGGAATGGTTAGCT
TACACCGCCAAAATGCTGGCGTAGTAGGTCAAGACTCTGAGACTATCGAACTC
GCACCTGAATACGCTGAGGCTATCGCAACCCGTGCAGGTGCGCTGGCTGGCAT
CTCTCCGATGTTCCAACCTTGCGTAGTTCCTCCTAAGCCGTGGACTGGCATTAC
TGGTGGTGGCTATTGGGCTAACGGTCGTCGTCCTCTGGCGCTGGTGCGTACTCA
CAGTAAGAAAGCACTGATGCGCTACGAAGACGTTTACATGCCTGAGGTGTACA
AAGCGATTAACATTGCGCAAAACACCGCATGGAAAATCAACAAGAAAGTCCTA
GCGGTCGCCAACGTAATCACCAAGTGGAAGCATTGTCCGGTCGAGGACATCCC
TGCGATTGAGCGTGAAGAACTCCCGATGAAACCGGAAGACATCGACATGAATC
CTGAGGCTCTCACCGCGTGGAAACGTGCTGCCGCTGCTGTGTACCGCAAGGAC
AAGGCTCGCAAGTCTCGCCGTATCAGCCTTGAGTTCATGCTTGAGCAAGCCAAT
AAGTTTGCTAACCATAAGGCCATCTGGTTCCCTTACAACATGGACTGGCGCGGT
CGTGTTTACGCTGTGTCAATGTTCAACCCGCAAGGTAACGATATGACCAAAGG
ACTGCTTACGCTGGCGAAAGGTAAACCAATCGGTAAGGAAGGTTACTACTGGC
TGAAAATCCACGGTGCAAACTGTGCGGGTGTCGATAAGGTTCCGTTCCCTGAG
CGCATCAAGTTCATTGAGGAAAACCACGAGAACATCATGGCTTGCGCTAAGTC
TCCACTGGAGAACACTTGGTGGGCTGAGCAAGATTCTCCGTTCTGCTTCCTTGC
GTTCTGCTTTGAGTACGCTGGGGTACAGCACCACGGCCTGAGCTATAACTGCTC
CCTTCCGCTGGCGTTTGACGGGTCTTGCTCTGGCATCCAGCACTTCTCCGCGAT
GCTCCGAGATGAGGTAGGTGGTCGCGCGGTTAACTTGCTTCCTggcggtTCTgga
ggtCACACTCTTTACGCCCCTGGAGGATACGACATTATGGGATATTTGGATCAG
ATTGGGAACCGCCCAAACCCTCAGGTCGAACTGGGGCCTGTGGACACGTCATG
TGCCCTGATCCTGTGCGATCTGAAGCAAAAGGACACTCCGATCGTCTACGCCT
CGGAAGCCTTCTTGTATATGACCGGATACAGCAATGCAGAGGTGCTCGGCAGG
AACTGCAGATTCCTGCAGTCCCCCGACGGGATGGTGAAACCAAAGTCGACTCG
CAAATATGTGGACTCGAACACGATCAACACCATCCGGAAGGCCATCGACCGG
AACGCCGAGGTCCAGGTGGAGGTGGTCAACTTTAAGAAGAACGGCCAGCGGT
TCGTGAACTTTCTGACCATCATTCCGGTCCGGGATGAAACCGGAGAGTACAGA
TACTCCATGGGATTCCAGTGCGAAACCGAATAACCTAGGAGGCATCaaataaaac
gaaaggctcggcgcgccactagtatgtacaagTAGTAATCTAGAGGGCCCTATTCTATAG

TGTCACCTAAATGCTAGAGCTCGCTGATCAGTCCGTCGACggatccaccggatctag
ataactgatcataatcagccataccacatttgtagaggttttacttgctttaaaaaacctcccacacctcccc
ctgaacctgaaacataaaatgaatgcaattgttgttgttaacttgtttattgcagcttataatggttacaaat
aaagcaatagcatcacaaatttcacaaataaagcatttttttcactgcattctagttgtggtttgtccaaact
catcaatgtatctta

2. mOptoT7 Version 2 (V2)

pMag codon optimized
T7pol (1-563)

ATGCACACCCTGTACgccccc GGCGGCTACGACATCATGGGCTACCTGCGCCAG
ATCCGCAACCGCCCCAACCCCCAGGTGGAGCTGGGCCCCGTGGACACCAGCT
GCGCCCTGATCCTGTGCGACCTGAAGCAGAAGGACACCCCCATCGTGTACGCC
AGCGAGGCCTTCCTGTACATGACCGGCTACAGCAACGCCGAGGTGCTGGGCC
GCAACTGCCGCTTCCTGCAGAGCCCCGACGGCATGGTGAAGCCCAAGAGCAC
CCGCAAGTACGTGGACAGCAACACCATCAACACCATGCGCAAGGCCATCGAC
CGCAACGCCGAGGTGCAGGTGGAGGTGGTGAACTTCAAGAAGAACGGCCAGC
GCTTCGTGAACTTCCTGACCATGATCCCCGTGCGCGACGAGACTGGCGAGTAC
CGCTACAGCATGGGCTTCCAGTGCGAGACTGAGggcggtTCTggaggtAGTGAAA
CCGTTCAGGACATCTACGGGATTGTTGCTAAGAAAGTCAACGAGATTCTACAA
GCAGACGCAATCAATGGGACCGATAACGAAGTAGTTACCGTGACCGATGAGA
ACACTGGTGAAATCTCTGAGAAAGTCAAGCTGGGCACTAAGGCACTGGCTGGT
CAATGGCTGGCTTACGGTGTTACTCGCAGTGTGACTAAGCGTTCAGTCATGACG
CTGGCTTACGGGTCCAAAGAGTTCGGCTTCCGTCAACAAGTGCTGGAAGATAC
CATTCAGCCAGCTATTGATTCCGGCAAGGGTCTGATGTTCACTCAGCCGAATCA
GGCTGCTGGATACATGGCTAAGCTGATTTGGGAATCTGTGAGCGTGACGGTGG

**TAGCTGCGGTTGAAGCAATGAACTGGCTTAAGTCTGCTGCTAAGCTGCTGGCT
GCTGAGGTCAAAGATAAGAAGACTGGAGAGATTCTTCGCAAGCGTTGCGCTGT
GCATTGGGTAACTCCTGATGGTTTCCCTGTGTGGCAGGAATACAAGAAGCCTAT
TCAGACGCGCTTGAACCTGATGTTCCTCGGTCAGTTCCGCTTACAGCCTACCAT
TAACACCAACAAAGATAGCGAGATTGATGCACACAAACAGGAGTCTGGTATCG
CTCCTAACTTTGTACACAGCCAAGACGGTAGCCACCTTCGTAAGACTGTAGTGT
GGGCACACGAGAAGTACGGAATCGAATCTTTTGCACTGATTCACGACTCCTTC
GGTACCATTCCGGCTGACGCTGCGAACCTGTTCAAAGCAGTGCGCGAAACTAT
GGTTGACACATATGAGTCTTGTGATGTACTGGCTGATTTCTACGACCAGTTCGC
TGACCAGTTGCACGAGTCTCAATTGGACAAAATGCCAGCACTTCCGGCTAAAG
GTAACTTGAACCTCCGTGACATCTTAGAGTcggacttcgcgtTCGCGTAA**

ATGAACACGATTAACatcgctaagaacgacTTCTCTGACATCGAACTGGCTGCTATC
CCGTTCAACACTCTGGCTGACCATTACGGTGAGCGTTTAGCTCGCGAACAGTTG
GCCCTTGAGCATGAGTCTTACGAGATGGGTGAAGCACGCTTCCGCAAGATGTT
TGAGCGTCAACTTAAAGCTGGTGAGGTTGCGGATAACGCTGCCGCCAAGCCTC
TCATCACTACCCTACTCCCTAAGATGATTGCACGCATCAACGACTGGTTTGAGG
AAGTGAAAGCTAAGCGCGGCAAGCGCCCGACAGCCTTCCAGTTCCTGCAAGA
AATCAAGCCGGAAGCCGTAGCGTACATCACCATTAAGACCACTCTGGCTTGCC
TAACCAGTGCTGACAATACAACCGTTCAGGCTGTAGCAAGCGCAATCGGTCGG
GCCATTGAGGACGAGGCTCGCTTCGGTCGTATCCGTGACCTTGAAGCTAAGCA
CTTCAAGAAAAACGTTGAGGAACAACTCAACAAGCGCGTAGGGCACGTCTACA
AGAAAGCATTTATGCAAGTTGTCGAGGCTGACATGCTCTCTAAGGGTCTACTCG
GTGGCGAGGCGTGGTCTTCGTGGCATAAGGAAGACTCTATTCATGTAGGAGTA
CGCTGCATCGAGATGCTCATTGAGTCAACCGGAATGGTTAGCTTACACCGCCA
AAATGCTGGCGTAGTAGGTCAAGACTCTGAGACTATCGAACTCGCACCTGAAT
ACGCTGAGGCTATCGCAACCCGTGCAGGTGCGCTGGCTGGCATCTCTCCGATG
TTCCAACCTTGCGTAGTTCCTCCTAAGCCGTGGACTGGCATTACTGGTGGTGGC
TATTGGGCTAACGGTCGTCGTCCTCTGGCGCTGGTGCGTACTCACAGTAAGAA
AGCACTGATGCGCTACGAAGACGTTTACATGCCTGAGGTGTACAAAGCGATTA

ACATTGCGCAAAACACCGCATGGAAAATCAACAAGAAAGTCCTAGCGGTCGCC
AACGTAATCACCAAGTGGAAGCATTGTCCGGTCGAGGACATCCCTGCGATTGA
GCGTGAAGAACTCCCGATGAAACCGGAAGACATCGACATGAATCCTGAGGCT
CTCACCGCGTGGAAACGTGCTGCCGCTGCTGTGTACCGCAAGGACAAGGCTCG
CAAGTCTCGCCGTATCAGCCTTGAGTTCATGCTTGAGCAAGCCAATAAGTTTGC
TAACCATAAGGCCATCTGGTTCCCTTACAACATGGACTGGCGCGGTCGTGTTTA
CGCTGTGTCAATGTTCAACCCGCAAGGTAACGATATGACCAAAGGACTGCTTA
CGCTGGCGAAAGGTAAACCAATCGGTAAGGAAGGTTACTACTGGCTGAAAATC
CACGGTGCAAACTGTGCGGGTGTCGATAAGGTTCCGTTCCCTGAGCGCATCAA
GTTCATTGAGGAAAACCACGAGAACATCATGGCTTGCGCTAAGTCTCCACTGG
AGAACACTTGGTGGGCTGAGCAAGATTCTCCGTTCTGCTTCCTTGCGTTCTGCTT
TGAGTACGCTGGGGTACAGCACCACGGCCTGAGCTATAACTGCTCCCTTCCGC
TGGCGTTTGACGGGTCTTGCTCTGGCATCCAGCACTTCTCCGCGATGCTCCGAG
ATGAGGTAGGTGGTCGCGCGGTTAACTTGCTTCCT

nMagHigh1 codon optimized
T7pol (1-563)

ATGAACACGATTAACatcgctaagaacgacTTCTCTGACATCGAACTGGCTGCTATC
CCGTTCAACACTCTGGCTGACCATTACGGTGAGCGTTTAGCTCGCGAACAGTTG
GCCCTTGAGCATGAGTCTTACGAGATGGGTGAAGCACGCTTCCGCAAGATGTT
TGAGCGTCAACTTAAAGCTGGTGAGGTTGCGGATAACGCTGCCGCCAAGCCTC
TCATCACTACCCTACTCCCTAAGATGATTGCACGCATCAACGACTGGTTTGAGG
AAGTGAAAGCTAAGCGCGGCAAGCGCCCGACAGCCTTCCAGTTCCTGCAAGA
AATCAAGCCGGAAGCCGTAGCGTACATCACCATTAAGACCACTCTGGCTTGCC
TAACCAGTGCTGACAATACAACCGTTCAGGCTGTAGCAAGCGCAATCGGTCGG
GCCATTGAGGACGAGGCTCGCTTCGGTCGTATCCGTGACCTTGAAGCTAAGCA
CTTCAAGAAAAACGTTGAGGAACAACTCAACAAGCGCGTAGGGCACGTCTACA
AGAAAGCATTTATGCAAGTTGTCGAGGCTGACATGCTCTCTAAGGGTCTACTCG
GTGGCGAGGCGTGGTCTTCGTGGCATAAGGAAGACTCTATTCATGTAGGAGTA
CGCTGCATCGAGATGCTCATTGAGTCAACCGGAATGGTTAGCTTACACCGCCA

AAATGCTGGCGTAGTAGGTCAAGACTCTGAGACTATCGAACTCGCACCTGAAT
ACGCTGAGGCTATCGCAACCCGTGCAGGTGCGCTGGCTGGCATCTCTCCGATG
TTCCAACCTTGCGTAGTTCCTCCTAAGCCGTGGACTGGCATTACTGGTGGTGGC
TATTGGGCTAACGGTCGTCGTCCTCTGGCGCTGGTGCGTACTCACAGTAAGAA
AGCACTGATGCGCTACGAAGACGTTTACATGCCTGAGGTGTACAAAGCGATTA
ACATTGCGCAAAACACCGCATGGAAAATCAACAAGAAAGTCCTAGCGGTCGCC
AACGTAATCACCAAGTGGAAGCATTGTCCGGTCGAGGACATCCCTGCGATTGA
GCGTGAAGAACTCCCGATGAAACCGGAAGACATCGACATGAATCCTGAGGCT
CTCACCGCGTGGAAACGTGCTGCCGCTGCTGTGTACCGCAAGGACAAGGCTCG
CAAGTCTCGCCGTATCAGCCTTGAGTTCATGCTTGAGCAAGCCAATAAGTTTGC
TAACCATAAGGCCATCTGGTTCCCTTACAACATGGACTGGCGCGGTCGTGTTTA
CGCTGTGTCAATGTTCAACCCGCAAGGTAACGATATGACCAAAGGACTGCTTA
CGCTGGCGAAAGGTAAACCAATCGGTAAGGAAGGTTACTACTGGCTGAAAATC
CACGGTGCAAACTGTGCGGGTGTCGATAAGGTTCCGTTCCCTGAGCGCATCAA
GTTCATTGAGGAAAACCACGAGAACATCATGGCTTGCGCTAAGTCTCCACTGG
AGAACACTTGGTGGGCTGAGCAAGATTCTCCGTTCTGCTTCCTTGCGTTCTGCTT
TGAGTACGCTGGGGTACAGCACCACGGCCTGAGCTATAACTGCTCCCTTCCGC
TGGCGTTTGACGGGTCTTGCTCTGGCATCCAGCACTTCTCCGCGATGCTCCGAG
ATGAGGTAGGTGGTCGCGCGGTTAACTTGCTTCCTggcggTTCTggaggtCACAC
CCTGTACGCCCCCGGCGGCTACGACATCATGGGCTACCTGGACCAGATCGGCA
ACCGCCCCAACCCCCAGGTGGAGCTGGGCCCCGTGGACACCAGCTGCGCCCT
GATCCTGTGCGACCTGAAGCAGAAGGACACCCCCATCGTGTACGCCAGCGAG
GCCTTCCTGTACATGACCGGCTACAGCAACGCCGAGGTGCTGGGCCGCAACTG
CCGCTTCCTGCAGAGCCCCGACGGCATGGTGAAGCCCAAGAGCACCCGCAAG
TACGTGGACAGCAACACCATCAACACCATCCGCAAGGCCATCGACCGCAACG
CCGAGGTGCAGGTGGAGGTGGTGAACTTCAAGAAGAACGGCCAGCGCTTCGT
GAACTTCCTGACCATCATCCCCGTGCGCGACGAGACGGGCGAGTACCGCTACA
GCATGGgcttcCAGTGCGAGACGGAGTAA

#### 3. mOptoT7 Version 3 (V3)

EF1a promoter
pMag Codon Optimized
T7pol (564-883)

#### SV40polyA

GCTCCGGTGCCCGTCAGTGGGCAGAGCGCACATCGCCCACAGTCCCCGAGAA
GTTGGGGGGAGGGGTCGGCAATTGAACCGGTGCCTAGAGAAGGTGGCGCGG
GGTAAACTGGGAAAGTGATGTCGTGTACTGGCTCCGCCTTTTTCCCGAGGGTG
GGGGAGAACCGTATATAAGTGCAGTAGTCGCCGTGAACGTTCTTTTTCGCAAC
GGGTTTGCCGCCAGAACACAGGTAAGTGCCGTGTGTGGTTCCCGCGGGCCTGG
CCTCTTTACGGGTTATGGCCCTTGCGTGCCTTGAATTACTTCCACGCCCCTGGCT
GCAGTACGTGATTCTTGATCCCGAGCTTCGGGTTGGAAGTGGGTGGGAGAGTT
CGAGGCCTTGCGCTTAAGGAGCCCCTTCGCCTCGTGCTTGAGTTGAGGCCTGG
CCTGGGCGCTGGGGCCGCCGCGTGCGAATCTGGTGGCACCTTCGCGCCTGTCT
CGCTGCTTTCGATAAGTCTCTAGCCATTTAAAATTTTTGATGACCTGCTGCGACG
CTTTTTTTCTGGCAAGATAGTCTTGTAAATGCGGGCCAAGATCTGCACACTGGT
ATTTCGGTTTTTGGGGCCGCGGGCGGCGACGGGGCCCGTGCGTCCCAGCGCA
CATGTTCGGCGAGGCGGGGCCTGCGAGCGCGGCCACCGAGAATCGGACGGG
GGTAGTCTCAAGCTGGCCGGCCTGCTCTGGTGCCTGGCCTCGCGCCGCCGTGT
ATCGCCCCGCCCTGGGCGGCAAGGCTGGCCCGGTCGGCACCAGTTGCGTGAG
CGGAAAGATGGCCGCTTCCCGGCCCTGCTGCAGGGAGCTCAAAATGGAGGAC
GCGGCGCTCGGGAGAGCGGGCGGGTGAGTCACCCACACAAAGGAAAAGGGC
CTTTCCGTCCTCAGCCGTCGCTTCATGTGACTCCACGGAGTACCGGGCGCCGTC
CAGGCACCTCGATTAGTTCTCGAGCTTTTGGAGTACGTCGTCTTTAGGTTGGGG
GGAGGGGTTTTATGCGATGGAGTTTCCCCACACTGAGTGGGTGGAGACTGAAG
TTAGGCCAGCTTGGCACTTGATGTAATTCTCCTTGGAATTTGCCCTTTTTGAGTTT
GGATCTTGGTTCATTCTCAAGCCTCAGACAGTGGTTCAAAGTTTTTTTCTTCCAT
TTCAGGTGTCGTGACATAAACTGCCCTAGAccacCTATGCACACCCTGTACGCCC
CCGGCGGCTACGACATCATGGGCTACCTGCGCCAGATCCGCAACCGCCCCAAC
CCCCAGGTGGAGCTGGGCCCCGTGGACACCAGCTGCGCCCTGATCCTGTGCG
ACCTGAAGCAGAAGGACACCCCCATCGTGTACGCCAGCGAGGCCTTCCTGTAC
ATGACCGGCTACAGCAACGCCGAGGTGCTGGGCCGCAACTGCCGCTTCCTGCA
GAGCCCCGACGGCATGGTGAAGCCCAAGAGCACCCGCAAGTACGTGGACAGC
AACACCATCAACACCATGCGCAAGGCCATCGACCGCAACGCCGAGGTGCAGG
TGGAGGTGGTGAACTTCAAGAAGAACGGCCAGCGCTTCGTGAACTTCCTGACC
ATGATCCCCGTGCGCGACGAGACTGGCGAGTACCGCTACAGCATGGGCTTCCA
GTGCGAGACTGAGggcggtTCTggaggtAGTGAAACCGTTCAGGACATCTACGGG

ATTGTTGCTAAGAAAGTCAACGAGATTCTACAAGCAGACGCAATCAATGGGAC
CGATAACGAAGTAGTTACCGTGACCGATGAGAACACTGGTGAAATCTCTGAGA
AAGTCAAGCTGGGCACTAAGGCACTGGCTGGTCAATGGCTGGCTTACGGTGTT
ACTCGCAGTGTGACTAAGCGTTCAGTCATGACGCTGGCTTACGGGTCCAAAGA
GTTCGGCTTCCGTCAACAAGTGCTGGAAGATACCATTCAGCCAGCTATTGATTC
CGGCAAGGGTCTGATGTTCACTCAGCCGAATCAGGCTGCTGGATACATGGCTA
AGCTGATTTGGGAATCTGTGAGCGTGACGGTGGTAGCTGCGGTTGAAGCAATG
AACTGGCTTAAGTCTGCTGCTAAGCTGCTGGCTGCTGAGGTCAAAGATAAGAA
GACTGGAGAGATTCTTCGCAAGCGTTGCGCTGTGCATTGGGTAACTCCTGATG
GTTTCCCTGTGTGGCAGGAATACAAGAAGCCTATTCAGACGCGCTTGAACCTG
ATGTTCCTCGGTCAGTTCCGCTTACAGCCTACCATTAACACCAACAAAGATAGC
GAGATTGATGCACACAAACAGGAGTCTGGTATCGCTCCTAACTTTGTACACAGC
CAAGACGGTAGCCACCTTCGTAAGACTGTAGTGTGGGCACACGAGAAGTACG
GAATCGAATCTTTTGCACTGATTCACGACTCCTTCGGTACCATTCCGGCTGACG
CTGCGAACCTGTTCAAAGCAGTGCGCGAAACTATGGTTGACACATATGAGTCTT
GTGATGTACTGGCTGATTTCTACGACCAGTTCGCTGACCAGTTGCACGAGTCTC
AATTGGACAAAATGCCAGCACTTCCGGCTAAAGGTAACTTGAACCTCCGTGAC
ATCTTAGAGTcggacttcgcgtTCGCGTAAATCCTAAACTAGTAAGTAGTAATCTAG
AGGGCCctattctatagtgtcacctaaatgctagagctcgctgatcagtccatagtcgacggatccacc
ggatctagataactgatcataatcagccataccacatttgtagaggttttacttgctttaaaaaacctcccac
acctccccctgaacctgaaacataaaatgaatgcaattgttgttgttaacttgtttattgcagcttataatgg
ttacaaataaagcaatagcatcacaaatttcacaaataaagcatttttttcactgcattctagttgtggtttgt
ccaaactcatcaatgtatctta

EF1a promoter
nMag High1 Codon Optimized
T7pol (564-883)
SV40polyA

GCTCCGGTGCCCGTCAGTGGGCAGAGCGCACATCGCCCACAGTCCCCGAGAA
GTTGGGGGGAGGGGTCGGCAATTGAACCGGTGCCTAGAGAAGGTGGCGCGG
GGTAAACTGGGAAAGTGATGTCGTGTACTGGCTCCGCCTTTTTCCCGAGGGTG
GGGGAGAACCGTATATAAGTGCAGTAGTCGCCGTGAACGTTCTTTTTCGCAAC
GGGTTTGCCGCCAGAACACAGGTAAGTGCCGTGTGTGGTTCCCGCGGGCCTGG
CCTCTTTACGGGTTATGGCCCTTGCGTGCCTTGAATTACTTCCACGCCCCTGGCT
GCAGTACGTGATTCTTGATCCCGAGCTTCGGGTTGGAAGTGGGTGGGAGAGTT
CGAGGCCTTGCGCTTAAGGAGCCCCTTCGCCTCGTGCTTGAGTTGAGGCCTGG
CCTGGGCGCTGGGGCCGCCGCGTGCGAATCTGGTGGCACCTTCGCGCCTGTCT
CGCTGCTTTCGATAAGTCTCTAGCCATTTAAAATTTTTGATGACCTGCTGCGACG
CTTTTTTTCTGGCAAGATAGTCTTGTAAATGCGGGCCAAGATCTGCACACTGGT
ATTTCGGTTTTTGGGGCCGCGGGCGGCGACGGGGCCCGTGCGTCCCAGCGCA
CATGTTCGGCGAGGCGGGGCCTGCGAGCGCGGCCACCGAGAATCGGACGGG
GGTAGTCTCAAGCTGGCCGGCCTGCTCTGGTGCCTGGCCTCGCGCCGCCGTGT
ATCGCCCCGCCCTGGGCGGCAAGGCTGGCCCGGTCGGCACCAGTTGCGTGAG
CGGAAAGATGGCCGCTTCCCGGCCCTGCTGCAGGGAGCTCAAAATGGAGGAC
GCGGCGCTCGGGAGAGCGGGCGGGTGAGTCACCCACACAAAGGAAAAGGGC
CTTTCCGTCCTCAGCCGTCGCTTCATGTGACTCCACGGAGTACCGGGCGCCGTC
CAGGCACCTCGATTAGTTCTCGAGCTTTTGGAGTACGTCGTCTTTAGGTTGGGG
GGAGGGGTTTTATGCGATGGAGTTTCCCCACACTGAGTGGGTGGAGACTGAAG
TTAGGCCAGCTTGGCACTTGATGTAATTCTCCTTGGAATTTGCCCTTTTTGAGTTT
GGATCTTGGTTCATTCTCAAGCCTCAGACAGTGGTTCAAAGTTTTTTTCTTCCAT
TTCAGGTGTCGTGACATAAACTGCCCTAGAccacCTATGAACACGATTAACATCG
CTAAGAACGACTTCTCTGACATCGAACTGGCTGCTATCCCGTTCAACACTCTGG
CTGACCATTACGGTGAGCGTTTAGCTCGCGAACAGTTGGCCCTTGAGCATGAG
TCTTACGAGATGGGTGAAGCACGCTTCCGCAAGATGTTTGAGCGTCAACTTAAA
GCTGGTGAGGTTGCGGATAACGCTGCCGCCAAGCCTCTCATCACTACCCTACTC
CCTAAGATGATTGCACGCATCAACGACTGGTTTGAGGAAGTGAAAGCTAAGCG
CGGCAAGCGCCCGACAGCCTTCCAGTTCCTGCAAGAAATCAAGCCGGAAGCC
GTAGCGTACATCACCATTAAGACCACTCTGGCTTGCCTAACCAGTGCTGACAAT
ACAACCGTTCAGGCTGTAGCAAGCGCAATCGGTCGGGCCATTGAGGACGAGG
CTCGCTTCGGTCGTATCCGTGACCTTGAAGCTAAGCACTTCAAGAAAAACGTTG
AGGAACAACTCAACAAGCGCGTAGGGCACGTCTACAAGAAAGCATTTATGCAA
GTTGTCGAGGCTGACATGCTCTCTAAGGGTCTACTCGGTGGCGAGGCGTGGTC

TTCGTGGCATAAGGAAGACTCTATTCATGTAGGAGTACGCTGCATCGAGATGCT
CATTGAGTCAACCGGAATGGTTAGCTTACACCGCCAAAATGCTGGCGTAGTAG
GTCAAGACTCTGAGACTATCGAACTCGCACCTGAATACGCTGAGGCTATCGCA
ACCCGTGCAGGTGCGCTGGCTGGCATCTCTCCGATGTTCCAACCTTGCGTAGTT
CCTCCTAAGCCGTGGACTGGCATTACTGGTGGTGGCTATTGGGCTAACGGTCGT
CGTCCTCTGGCGCTGGTGCGTACTCACAGTAAGAAAGCACTGATGCGCTACGA
AGACGTTTACATGCCTGAGGTGTACAAAGCGATTAACATTGCGCAAAACACCG
CATGGAAAATCAACAAGAAAGTCCTAGCGGTCGCCAACGTAATCACCAAGTGG
AAGCATTGTCCGGTCGAGGACATCCCTGCGATTGAGCGTGAAGAACTCCCGAT
GAAACCGGAAGACATCGACATGAATCCTGAGGCTCTCACCGCGTGGAAACGT
GCTGCCGCTGCTGTGTACCGCAAGGACAAGGCTCGCAAGTCTCGCCGTATCAG
CCTTGAGTTCATGCTTGAGCAAGCCAATAAGTTTGCTAACCATAAGGCCATCTG
GTTCCCTTACAACATGGACTGGCGCGGTCGTGTTTACGCTGTGTCAATGTTCAA
CCCGCAAGGTAACGATATGACCAAAGGACTGCTTACGCTGGCGAAAGGTAAA
CCAATCGGTAAGGAAGGTTACTACTGGCTGAAAATCCACGGTGCAAACTGTGC
GGGTGTCGATAAGGTTCCGTTCCCTGAGCGCATCAAGTTCATTGAGGAAAACC
ACGAGAACATCATGGCTTGCGCTAAGTCTCCACTGGAGAACACTTGGTGGGCT
GAGCAAGATTCTCCGTTCTGCTTCCTTGCGTTCTGCTTTGAGTACGCTGGGGTAC
AGCACCACGGCCTGAGCTATAACTGCTCCCTTCCGCTGGCGTTTGACGGGTCTT
GCTCTGGCATCCAGCACTTCTCCGCGATGCTCCGAGATGAGGTAGGTGGTCGC
GCGGTTAACTTGCTTCCTggcggTTCTggaggtCACACCCTGTACGCCCCCGGCG
GCTACGACATCATGGGCTACCTGGACCAGATCGGCAACCGCCCCAACCCCCAG
GTGGAGCTGGGCCCCGTGGACACCAGCTGCGCCCTGATCCTGTGCGACCTGAA
GCAGAAGGACACCCCCATCGTGTACGCCAGCGAGGCCTTCCTGTACATGACCG
GCTACAGCAACGCCGAGGTGCTGGGCCGCAACTGCCGCTTCCTGCAGAGCCC
CGACGGCATGGTGAAGCCCAAGAGCACCCGCAAGTACGTGGACAGCAACACC
ATCAACACCATCCGCAAGGCCATCGACCGCAACGCCGAGGTGCAGGTGGAGG
TGGTGAACTTCAAGAAGAACGGCCAGCGCTTCGTGAACTTCCTGACCATCATC
CCCGTGCGCGACGAGACGGGCGAGTACCGCTACAGCATGGGCTTCCAGTGCG
AGACGGAGGGATCCTAAACTAGTAAGTAGTAATCTAGAGGGCCctattctatagtgt
cacctaaatgctagagctcgctgatcagtccatagtcgacggatccaccggatctagataactgatcata
atcagccataccacatttgtagaggttttacttgctttaaaaaacctcccacacctccccctgaacctgaaa
cataaaatgaatgcaattgttgttgttaacttgtttattgcagcttataatggttacaaataaagcaatagca

tcacaaatttcacaaataaagcatttttttcactgcattctagttgtggtttgtccaaactcatcaatgtatctt
a

4. mOptoT7 reporter

T7 promoter
IRES2
mRuy3
T7 terminator

taatacgactcactatagggagacccaagcttatgcatgcggccgcatctagagggcccggatccg
cccctctccctcccccccccctaacgttactggccgaagccgcttggaataaggccggtgtgcgtttgtct
atatgttattttccaccatattgccgtcttttggcaatgtgagggcccggaaacctggccctgtcttcttgac
gagcattcctaggggtctttcccctctcgccaaaggaatgcaaggtctgttgaatgtcgtgaaggaagc
agttcctctggaagcttcttgaagacaaacaacgtctgtagcgaccctttgcaggcagcggaacccccc
acctggcgacaggtgcctctgcggccaaaagccacgtgtataagatacacctgcaaaggcggcacaa
ccccagtgccacgttgtgagttggatagttgtggaaagagtcaaatggctctcctcaagcgtattcaaca
aggggctgaaggatgcccagaaggtaccccattgtatgggatctgatctggggcctcggtacacatgc
tttacatgtgtttagtcgaggttaaaaaaacgtctaggccccccgaaccacggggacgtggttttcctttg
aaaaacacgatgataatatggccacaaccATGGTGTCTAaGGGCGAAGAGctGATCAAGG
AAAATATGCGTATGAAGGTGGTCATGGAAGGTTCGGTCAACGGCCACCAATTC
AAATGCACAGGTGAAGGAGAAGGCAGACCGTACGAGGGAGTGCAAACCATG
AGGATCAAAGTCATCGAGGGAGGACCCCTGCCATTTGCCTTTGACATTCTTGCC
ACGTCGTTCATGTATGGCAGCCGTACCTTTATCAAGTACCCGGCCGACATCCCT
GATTTCTTTAAACAGTCCTTTCCTGAGGGTTTTACTTGGGAAAGAGTTACGAGAT
ACGAAGATGGTGGAGTCGTCACCGTCACGCAGGACACCAGCCTTGAGGATGG
CGAGCTCGTCTACAACGTCAAGGTCAGAGGGGTAAACTTTCCCTCCAATGGTC
CCGTGATGCAGAAGAAGACCAAGGGTTGGGAGCCTAATACAGAGATGATGTA
TCCAGCAGATGGTGGTCTGAGAGGATACACTGACATCGCACTGAAAGTTGATG
GTGGTGGCCATCTGCACTGCAACTTCGTGACAACTTACAGGTCAAAAAAGACC
GTCGGGAACATCAAGATGCCCGGTGTCCATGCCGTTGATCACCGCCTGGAAAG
GATCGAGGAGAGTGACAATGAAACCTACGTAGTGCAAAGAGAAGTGGCAGTT
GCCAAATACAGCAACCTTGGTGGTGGCATGGACGAGCTgtacaagtaaagcgctgaa
ggtgaatggcagctggttctgcatgtttgggctaaagttgaagctgacgtcgctggtcatggtcaggac

atcttgattcgactgttcaaatctcatccggaaactctggaaaaattcgatcgtttcaaacatctgaaaact
gaagctgaaatgaaagcttctg
atagcataaccccttggggcctctaaacgggtcttgaggggttttttg

#### 5. mOptoT7 reporter with polyA

T7 promoter
IRES2
mRuy3
polyA
T7 terminator

taatacgactcactatagggagacccaagcttatgcatgcggccgcatctagagggcccggatccg
cccctctccctcccccccccctaacgttactggccgaagccgcttggaataaggccggtgtgcgtttgtct
atatgttattttccaccatattgccgtcttttggcaatgtgagggcccggaaacctggccctgtcttcttgac
gagcattcctaggggtctttcccctctcgccaaaggaatgcaaggtctgttgaatgtcgtgaaggaagc
agttcctctggaagcttcttgaagacaaacaacgtctgtagcgaccctttgcaggcagcggaacccccc
acctggcgacaggtgcctctgcggccaaaagccacgtgtataagatacacctgcaaaggcggcacaa
ccccagtgccacgttgtgagttggatagttgtggaaagagtcaaatggctctcctcaagcgtattcaaca
aggggctgaaggatgcccagaaggtaccccattgtatgggatctgatctggggcctcggtacacatgc
tttacatgtgtttagtcgaggttaaaaaacgtctaggccccccgaaccacggggacgtggttttcctttg
aaaaacacgatgataatatggccacaaccATGGTGTCTAaGGGCGAAGAGctGATCAAGG
AAAATATGCGTATGAAGGTGGTCATGGAAGGTTCGGTCAACGGCCACCAATTC
AAATGCACAGGTGAAGGAGAAGGCAGACCGTACGAGGGAGTGCAAACCATG
AGGATCAAAGTCATCGAGGGAGGACCCCTGCCATTTGCCTTTGACATTCTTGCC
ACGTCGTTCATGTATGGCAGCCGTACCTTTATCAAGTACCCGGCCGACATCCCT
GATTTCTTTAAACAGTCCTTTCCTGAGGGTTTTACTTGGGAAAGAGTTACGAGAT
ACGAAGATGGTGGAGTCGTCACCGTCACGCAGGACACCAGCCTTGAGGATGG
CGAGCTCGTCTACAACGTCAAGGTCAGAGGGGTAAACTTTCCCTCCAATGGTC
CCGTGATGCAGAAGAAGACCAAGGGTTGGGAGCCTAATACAGAGATGATGTA
TCCAGCAGATGGTGGTCTGAGAGGATACACTGACATCGCACTGAAAGTTGATG
GTGGTGGCCATCTGCACTGCAACTTCGTGACAACTTACAGGTCAAAAAAGACC
GTCGGGAACATCAAGATGCCCGGTGTCCATGCCGTTGATCACCGCCTGGAAAG
GATCGAGGAGAGTGACAATGAAACCTACGTAGTGCAAAGAGAAGTGGCAGTT

GCCAAATACAGCAACCTTGGTGGTGGCATGGACGAGCTgtacaagtaaTTCTagct
gtacaagTAAAcgcgtcttaaataaaaaaaaaaaaaaaaaaaaaaaaaaaaaaaaaaaaaaaaaaaa
aaaaaaaaaaaaaaaaaaaaaaaaaaaaaaaaaaaaaaaaaaaaaaaaaaaaaaaaaaagcgatc
gcgggcGGCCGCAGGTACCCTCGAGTCATCCtaaagcgctgaaggtgaatggcagctggt
tctgcatgtttgggctaaagttgaagctgacgtcgctggtcatggtcaggacatcttgattcgactgttca
aatctcatccggaaactctggaaaaattcgatcgtttcaaacatctgaaaactgaagctgaaatgaaagc
ttctgatagcataaccccttggggcctctaaacgggtcttgaggggttttttg

##### 6. mOptoT7 reporter with 8XPepper RNA aptamer

T7 promoter
8XPepper
T7 terminator

TAATACGACTCACTATAGGGAGTATGCTTGGTACCGAGCTCGGATCcACTAGTC
CAGTGTGGTGGAATTCCCCACGGAGGATCCCCAATCGTGGCGTGTCGGCCTCT
CCCAATCGTGGCGTGTCGGCCTCTCCCAATCGTGGCGTGTCGGCCTCTCCCAAT
CGTGGCGTGTCGGCCTCTCCCAATCGTGGCGTGTCGGCCTCTCCCAATCGTGG
CGTGTCGGCCTCTCCCAATCGTGGCGTGTCGGCCTCTCCCAATCGTGGCGTGTC
GGCCTCTCTTCGGAGAGGCACTGGCGCCGGAGAGGCACTGGCGCCGGAGAGG
CACTGGCGCCGGAGAGGCACTGGCGCCGGAGAGGCACTGGCGCCGGAGAGG
CACTGGCGCCGGAGAGGCACTGGCGCCGGAGAGGCACTGGCGCCGGGATCCT
CCGTGGGCTCGAGTCTAGAGGgCCCGTTTAAACCCGCTGATCAATCCtaaagcgc
tgaaggtgaatggcagctggttctgcatgtttgggctaaagttgaagctgacgtcgctggtcatggtca
ggacatcttgattcgactgttcaaatctcatccggaaactctggaaaaattcgatcgtttcaaacatctgaa
aactgaagctgaaatgaaagcttctgatagcataaccccttggggcctctaaacgggtcttgaggggttt
tttg

##### 7. mOptoT7 reporter with shRNA

T7 promoter
shRNA
T7 terminator

TAATACGACTCACTATAGGGAGTATGGCAAGCTGACCCTGAAGTTCATTCAAG
AGATGAACTTCAGGGTCAGCTTGCATCCtaaagcgctgaaggtgaatggcagctggttct
gcatgtttgggctaaagttgaagctgacgtcgctggtcatggtcaggacatcttgattcgactgttcaaat
ctcatccggaaactctggaaaaattcgatcgtttcaaacatctgaaaactgaagctgaaatgaaagcttct
gatagcataaccccttggggcctctaaacgggtcttgaggggttttttg
